# Vision-language encoding models reveal an image-computable food-quality dimension in human occipitotemporal cortex

**DOI:** 10.64898/2026.08.25.747006

**Authors:** Giuseppe Marrazzo, Anne Roefs, Leonardo Pimpini

## Abstract

Perceived calorie content contributes to neural representational structure in human ventral visual cortex, yet it remains unclear whether this reflects an abstract nutritional signal or whether perceived calorie is largely recoverable from the visual-semantic structure of the food image itself. In 25 female participants who passively viewed 96 food images during functional MRI, we decomposed perceived calorie ratings into a component predicted from CLIP (Contrastive Language-Image Pretraining) image embeddings, a vision-language model that captures high-level visual-semantic image structure, and a residual component not captured by this CLIP-based prediction. We then tested their respective contributions to neural prediction using cross-validated banded ridge encoding models. The CLIP-predictable component organized foods along a processedness and naturalness dimension, separating raw single-ingredient foods from prepared and energy-dense foods. Adding this component to a visual-semantic baseline improved neural prediction progressively along the ventral visual hierarchy, with the strongest relative contribution in higher-level ventral temporal cortex. These findings indicate that calorie-related encoding in ventral visual cortex is carried mainly by a shared food-quality axis indexing processedness, naturalness, and perceived healthiness, a substantial part of which is recoverable from image-computable visual-semantic structure, rather than providing evidence for an isolated abstract representation of caloric magnitude. Because these results derive from a reanalysis of 25 female participants viewing a fixed set of 96 images, generalization to broader populations and larger, more varied stimulus sets remains to be established.

## Introduction

The human visual system does not represent food images as collections of pixels. By the time visual information reaches higher-level ventral temporal cortex, neural responses are organized by object category, semantic content, and behaviorally relevant stimulus properties rather than by low-level image statistics alone(DiCarlo, Zoccolan, & Rust, 2012; Grill-Spector & Weiner, 2014). Food stimuli are a particularly informative test case for this kind of high-level visual organization because they vary simultaneously in visual appearance, category membership, and subjective attributes such as perceived caloric content, palatability, and familiarity (Blechert, Lender, Polk, Busch, & Ohla, 2019; Blechert, Meule, Busch, & Ohla, 2014; Rangel, 2013; van der Laan, de Ridder, Viergever, & Smeets, 2011). These dimensions may be partially overlapping rather than independently represented in visual cortex, making food images well suited for testing how visual, semantic, and subjective information are organized in ventral visual representations. Consistent with this view, recent multivariate and data-driven neuroimaging studies have shown that food representations in ventral visual cortex reflect category-selective and food-relevant structure that cannot be reduced to low-level visual features alone (Jain et al., 2023; Khosla, Ratan Murty, & Kanwisher, 2022; Moerel, Psihoyos, & Carlson, 2024; Pennock et al., 2023).

In a companion study to the present work (Marrazzo et al., 2026), we found that food-related representational structure in lateral and ventral occipitotemporal cortex was not fully explained by the tested low- and high-level visual feature models. That study showed that perceived caloric content and perceived health value were strongly overlapping in representational space, and did not provide evidence for a separable perceived-calorie code distinct from this shared subjective food-property axis. Thus, the companion RSA analysis established where food-related representational structure was detectable and how it related to subjective and categorical food dimensions, but it could not determine what kind of information gave rise to the shared axis. In particular, it remained unclear whether this axis was primarily recoverable from the visual-semantic structure of the food image itself, or whether it reflected calorie-related information not captured by the tested image-computable representations. The present study addresses this unresolved question by re-analysing the same dataset with encoding models, asking what the shared food-quality axis is made of and how its neural relevance changes along the ventral visual hierarchy.

This distinction is important because a positive association between perceived calorie and neural responses is ambiguous with respect to mechanism. Under one interpretation, the neural effect reflects a representation of caloric magnitude itself: an abstract nutritional signal that may guide food valuation and intake regulation, consistent with evidence that reward-related regions can track caloric content implicitly (Tang, Fellows, & Dagher, 2014). Under another interpretation, perceived calorie is partly recoverable from the visual-semantic structure of the food image itself, that is, from what can be inferred from how the food appears and from the semantic associations carried by that appearance. These accounts are not a contrast between learned and innate information, since all high-level visual representation is learned from experience. Rather, the relevant distinction is whether the neural-predictive component of perceived calorie is mainly captured by image-computable visual-semantic structure, or whether it depends on residual calorie-related information not captured by the tested image-computable models. Consistent with this possibility, perceived calorie and perceived health value show strong shared structure in food-image datasets (Carels, Konrad, & Harper, 2007; Charbonnier, van Meer, van der Laan, Viergever, & Smeets, 2016), matching the shared subjective axis identified in the companion RSA analysis of the present stimulus set. Thus, perceived calorie may index a broad food-quality dimension rather than caloric magnitude specifically. Distinguishing these possibilities requires decomposing perceived calorie into the component recoverable from visual-semantic image structure and the residual component not captured by the tested visual-semantic model, and testing their respective contributions to neural prediction.

Recent advances in vision-language models provide a principled basis for this decomposition. CLIP (Contrastive Language-Image Pretraining) (Radford et al., 2021), trained on large-scale image-text pairs using contrastive learning, captures high-level visual-semantic image structure that goes beyond object recognition, including conceptual and categorical properties of visual scenes. Critically, CLIP embeddings encode not just what is in an image but how it relates to the broader space of linguistic and perceptual concepts associated with it (Radford et al., 2021), making them well-suited for characterizing the visual-semantic content of perceived calorie and, by extension, the broader food-quality dimension it indexes. We therefore used CLIP embeddings to predict perceived calorie ratings across food images. The predicted values, which we refer to as CaloriePredCLIP, capture the part of perceived calorie that is recoverable from the visual-semantic image structure captured by CLIP. The residual values, which we refer to as CalorieResCLIP, capture the remaining variance in perceived calorie ratings after removing the CLIP-predictable component. Because CLIP is a single vision-language model with a specific architecture, training history, and representational format, this residual should be interpreted as variance not recovered by the tested CLIP representation, rather than as definitive evidence for information that is unavailable from images in principle. If the neural-relevant component of perceived calorie is primarily the part recoverable from the tested visual-semantic image representation, CaloriePredCLIP should carry most of the calorie-related neural-predictive information. If, by contrast, neural responses reflect calorie-related information beyond the tested visual-semantic representation, CalorieResCLIP should carry independent neural-predictive information.

The present study directly tests this question using encoding models, which predict measured neural responses from stimulus feature representations and evaluate prediction accuracy on held-out data. This framework provides a principled basis for adjudicating between competing hypotheses of what information a brain region represents (Naselaris, Kay, Nishimoto, & Gallant, 2011; Yamins & DiCarlo, 2016): by specifying feature spaces that operationalize alternative hypotheses and comparing their held-out predictive performance in a nested model family, encoding models allow researchers to test not merely whether a brain region responds to a stimulus dimension, but whether its response reflects a specific type of information, image-computable structure, semantic content, or behavioral ratings, and where along the cortical hierarchy that information is most prominently encoded (Huth, Nishimoto, Vu, & Gallant, 2012; Kay, Naselaris, Prenger, & Gallant, 2008; Naselaris et al., 2011). This approach has proven productive for characterizing the representational content of visual cortex across domains ranging from low-level image statistics to high-level semantic categories (Mitchell et al., 2008) (Marrazzo, De Martino, Lage-Castellanos, Vaessen, & de Gelder, 2023; Marrazzo, De Martino, Mukovskiy, Giese, & de Gelder, 2025; Nishimoto et al., 2011).

Here, we re-analysed the fMRI dataset reported in Marrazzo et al.(Marrazzo et al., 2026) using a nested family of cross-validated banded ridge encoding models(Nunez-Elizalde, Huth, & Gallant, 2019). The critical test was whether a targeted CLIP-derived axis predicting perceived calorie could explain neural responses beyond a visual baseline spanning low-level image features, intermediate visual representations, and high-level deep-network activations, including CLIP itself, with complementary analyses testing baselines in which CLIP was either excluded or modeled as a separately regularized feature band. This makes the analysis distinct from simply asking whether CLIP predicts visual cortex: we asked whether the part of perceived calorie that is expressible within CLIP’s visual-semantic image space captures a food-quality dimension with additional neural-predictive value. We predicted that if calorie-related neural prediction primarily reflects visual-semantic food structure rather than residual perceived-calorie variance, CaloriePredCLIP should outperform CalorieResCLIP, with the strongest effect in higher-level ventral temporal cortex, where semantic food representations are expected to be most prominent.

## Results

### Perceived calorie ratings were strongly predictable from CLIP image embeddings

At the group level, CLIP was the strongest predictor of perceived calorie, explaining 76.9% of cross-validated variance and showing a strong association with perceived calorie ratings (Pearson r = .878, Spearman ρ = .869; Figure 1B; Supplementary Table S1 and Figure S1). By contrast, LowVis features carried essentially no predictive information about perceived calorie, whereas HighVis features showed only moderate prediction (Supplementary Table S1). The same pattern was present at the individual-rating level, where CLIP predicted participants’ perceived calorie ratings with a mean CV R2 of .535 (Supplementary Figure S1 and Table S2).

**Figure 1.**
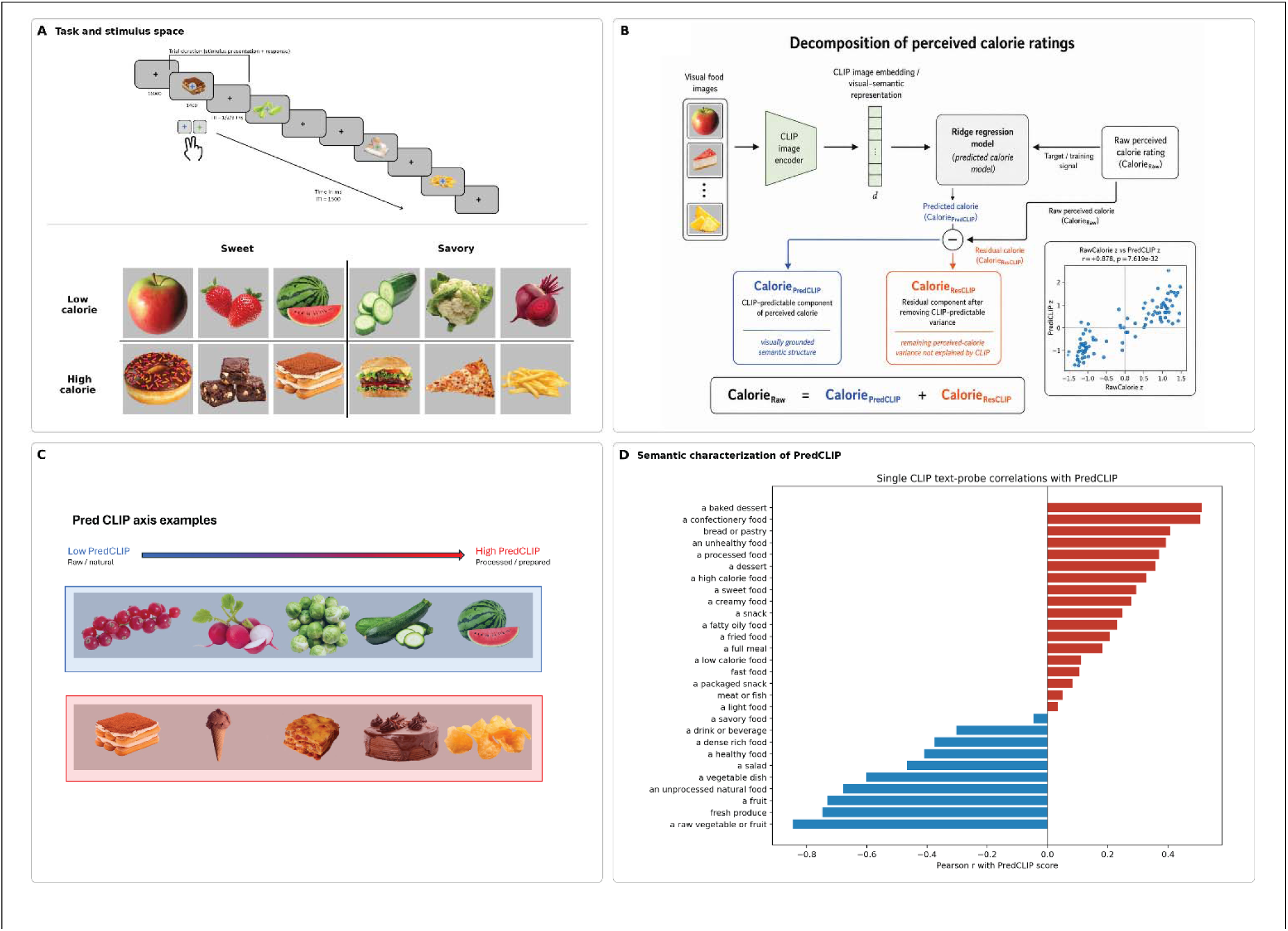
Experimental design and characterization of the CLIP-predicted perceived-calorie axis. (A) Task and stimulus space as in (Marrazzo et al., 2026). The stimulus set consisted of 96 food images arranged in a balanced 2 × 2 design crossing perceived calorie content, low versus high, with taste category, sweet versus savory. During fMRI, participants viewed the food images in a rapid event-related design while performing an orthogonal fixation-color discrimination task, ensuring attention to the stimuli without requiring explicit food evaluation. In Supplementary Figure S5 we show the complete set of 96 food images used in the fMRI experiment. (B) Schematic of the CLIP-based decomposition of perceived calorie ratings into CaloriePredCLIP and CalorieResCLIP. The inset scatterplot shows the strong association between perceived calorie and the CLIP-predicted component. (C) Representative images from the low and high ends of the PredCLIP axis. (D) Semantic characterization of the PredCLIP axis using CLIP text-probe correlations. Together, these analyses indicate that the CLIP-predicted calorie component primarily reflects an image-computable processedness/naturalness axis rather than an abstract nutritional calorie signal.

This result is central to the interpretation of the neural models. The aim of the decomposition was not to show that CaloriePredCLIP contains information independent of CLIP. Rather, CaloriePredCLIP was designed to isolate the specific direction within CLIP-readable visual-semantic space that best captures perceived calorie. The fact that CLIP predicted perceived calorie substantially better than LowVis or the broader HighVis composite indicates that perceived calorie is embedded in high-level visual-semantic structure, while not being reducible to low-level image statistics. Thus, the subsequent neural analyses test whether this specific calorie-aligned semantic direction is neurally relevant, not whether CLIP-like semantics are absent from the visual baseline.

Inspection of the CaloriePredCLIP axis showed that it separated raw, natural, minimally prepared foods from processed, prepared, dessert-like, and energy-dense foods (Figure 1C). Text-probe analyses confirmed this interpretation: CaloriePredCLIP was positively associated with processed, prepared, pastry, confectionery, and high-calorie descriptors, and negatively associated with raw, natural, fruit, and vegetable descriptors (Figure 1D). Thus, the CaloriePredCLIP axis was characterized by an interpretable processedness and naturalness contrast recoverable from image structure. Additional feature-space and representational-diagnostic analyses are reported in Supplementary Figure S1 and Supplementary Tables S1-S3.

A complementary layerwise analysis showed that processing/preparation structure was expressed most strongly in later CLIP representations rather than in early layers (Supplementary Figure S2), further supporting the interpretation that CaloriePredCLIP captured higher-level visual-semantic structure.

### CaloriePredCLIP improved prediction beyond M0 and increased along the visual hierarchy

M0 produced reliable held-out prediction in EarlyVisual cortex (mean r_joint = .048, 95% CI [.037, .058], q < .001) and IntermediateVisual cortex (mean r_joint = .069, 95% CI [.053, .086], q < .001), but was not reliably above zero in HighLevelVTC (mean r_joint = .002, 95% CI [-.006, .010], q = .283; Figure 2A; Supplementary Table S4). The weak performance of M0 in HighLevelVTC should not be interpreted as evidence that high-level visual-semantic information is absent from this region. M0 used a broad HighVis band in which AlexNet, CORnet, and CLIP features were combined and compressed into a single multivariate visual-semantic baseline. Therefore, poor M0 performance in HighLevelVTC may reflect the difficulty of recovering a specific food-semantic axis from a broad, regularized, high-dimensional feature space. The relevant question is whether the calorie-aligned direction isolated from CLIP captures neural-relevant structure that is not efficiently recovered by this broader baseline.

**Figure 2.**
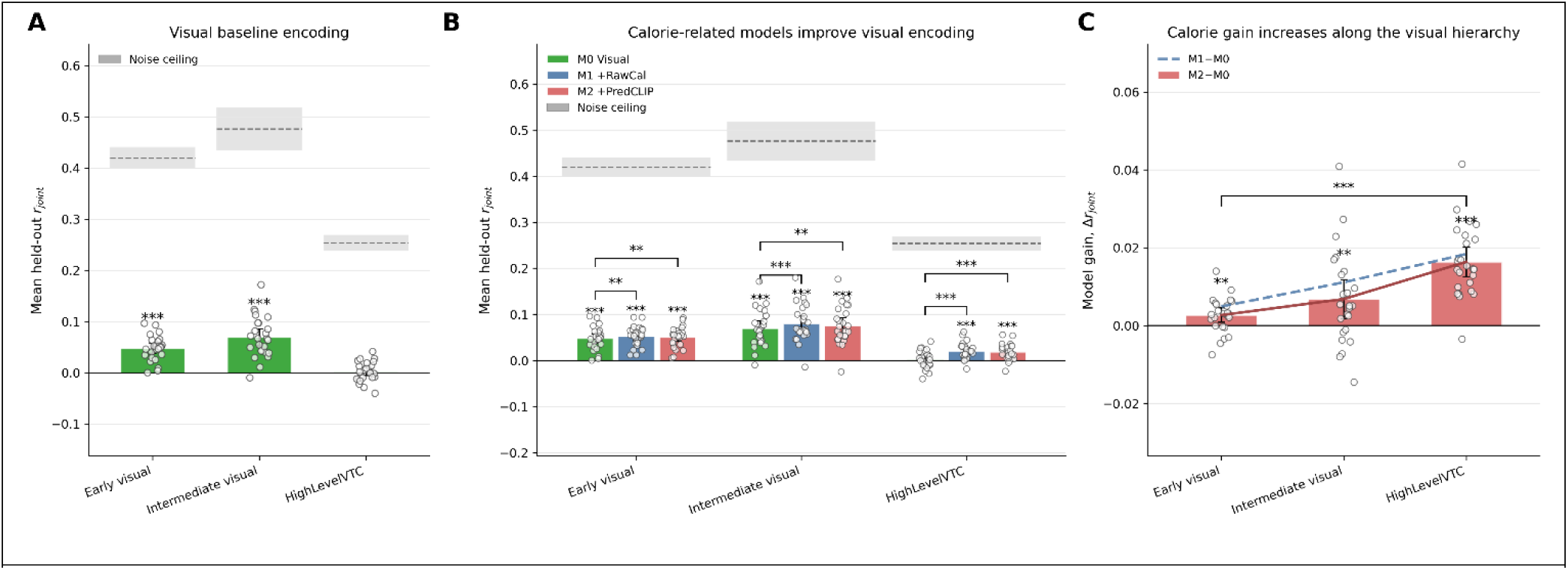
CaloriePredCLIP improves neural prediction increasingly along the visual hierarchy. (A) Held-out prediction accuracy for the visual-semantic baseline model, M0, across EarlyVisual, IntermediateVisual, and HighLevelVTC. Green bars show group-mean cross-validated prediction accuracy (r_joint) for M0, with error bars indicating 95% confidence intervals across participants. Gray horizontal bands indicate split-half noise-ceiling estimates for each ROI, shown as the mean with 95% confidence interval. Noise ceilings were computed for each participant by repeatedly splitting the available stimulus repetitions into two approximately equal halves, estimating condition-wise responses independently in each half, and applying the Spearman–Brown and correlation-scale corrections described in Methods (B) Prediction accuracy for M0, M1, and M2 across the three ROIs. M1 added perceived calorie to the visual-semantic baseline, whereas M2 added the CLIP-predictable component of perceived calorie. Both calorie-related models improved prediction relative to M0, with the largest gains observed in HighLevelVTC. Gray bands again indicate ROI-level split-half noise ceilings. (C) Model gains relative to M0, shown as Δr_joint for M1 − M0 and M2 − M0. Both gains were tested for a linear increase across the visual hierarchy. The primary theoretical contrast was M2 − M0, testing whether the CLIP-predictable component of perceived calorie contributed increasingly from EarlyVisual to HighLevelVTC; M1 − M0 is shown and tested as the corresponding perceived-calorie gain. In all panels, points represent individual participants, bars show group means, and error bars indicate 95% confidence intervals. Significance markers indicate FDR-corrected effects at q < .05. Asterisks above individual bars denote above-zero prediction or gain; brackets denote paired model comparisons.

Adding CaloriePredCLIP improved prediction beyond M0 in all three ROIs (Figure 2B-C). The M2-M0 gain was small in EarlyVisual cortex (Δr_joint = .003, 95% CI [.001, .005], q = .008), larger in IntermediateVisual cortex (Δr_joint = .007, 95% CI [.002, .012], q = .008), and largest in HighLevelVTC (Δr_joint = .016, 95% CI [.013, .020], q < .001; Supplementary Table S5). The significant gain in EarlyVisual cortex was not specific to the calorie-aligned ordering of stimuli, as confirmed by the label-shuffle specificity analysis reported below. The M2-M0 gain increased significantly along the visual hierarchy, with a mean slope of .007 Δr_joint per hierarchy step (q < .001). Pairwise hierarchy tests confirmed that the M2-M0 gain was larger in HighLevelVTC than in EarlyVisual (.014 Δr_joint; q < .001) cortex and IntermediateVisual cortex, whereas the IntermediateVisual > EarlyVisual contrast did not survive FDR correction (.004 Δr_joint; q = .057) (Supplementary Table S5d).

This hierarchical pattern is central to the main interpretation. If the CaloriePredCLIP effect were driven primarily by low-level visual confounds, the gain would be expected to peak in early visual cortex. Instead, the effect was strongest in HighLevelVTC, supporting the interpretation that CaloriePredCLIP captured higher-level visual-semantic structure.

The CLIP-separated diagnostic confirmed that weak baseline prediction in HighLevelVTC reflected joint feature compression rather than an absence of visual signal. Removing CLIP (D0) left HighLevelVTC prediction at chance (mean r = 0.000, not significant), whereas restoring CLIP as its own band (D1) raised it to r = 0.028 (D1 > D0: Δr = 0.028, 95% CI [0.023, 0.033], q < .001). Adding CaloriePredCLIP improved prediction further, specifically in HighLevelVTC (D2 > D1: Δr = 0.008, 95% CI [0.005, 0.011], q < .001), a ∼29% relative increase over the CLIP band. The same comparison was non-significant in EarlyVisual (Δr = 0.001, q = .22) and IntermediateVisual (Δr = 0.001, q = .37). This converges with the label-shuffle analysis: the EarlyVisual M2-M0 gain, which was not specific to the calorie-aligned stimulus ordering, also did not survive the CLIP-separated baseline, indicating it did not reflect calorie-aligned structure. The calorie-aligned gain over the visual baseline reported above is therefore not an artifact of an under-expressed CLIP representation: it persists against a CLIP band fit with its own regularization and is confined to higher ventral temporal cortex (Supplementary Figure S6; Supplementary Table S10).

### The CLIP-predictable component explained the calorie effect better than the CLIP-residual component

M1 and M2 showed closely matched prediction profiles: both improved prediction over M0 and both showed their largest gains in HighLevelVTC. M2 was numerically below M1 in all three ROIs (Supplementary Table S5), consistent with the expectation that CalorieRaw, which equals CaloriePredCLIP plus CalorieResCLIP by construction, will predict at least as well as either component alone. M2 did not outperform M1 in any ROI, consistent with the strong relationship between CalorieRaw and CaloriePredCLIP shown in Figure 1B. The decomposition therefore adds interpretive specificity rather than superior prediction over perceived calorie. This pattern is expected if perceived calorie ratings are dominated by a CLIP-readable semantic axis. Because CaloriePredCLIP is a supervised projection of CLIP onto perceived calorie, it should not be interpreted as a feature space independent of CLIP. Instead, its value is that it identifies which part of the visual-semantic food-image space is aligned with perceived calorie and tests whether that axis is neurally predictive. In this sense, the comparison between M2 and M3 is more informative than the comparison between M2 and M1: M2 and M1 ask whether the same dominant calorie-aligned axis predicts neural responses, whereas M2 versus M3 tests whether the neural-relevant component of perceived calorie is the CLIP-readable component or the residual component not captured by CLIP.

The key specificity result was that M2 outperformed M3 in IntermediateVisual cortex (Δr_joint = .014, 95% CI [.005, .023], q = .004) and HighLevelVTC (Δr_joint = .008, 95% CI [.004, .013], q = .001), but not in EarlyVisual cortex (Δr_joint = .000, 95% CI [-.004, .004], q = .871; Supplementary Table S5). Thus, the neural-relevant component of perceived calorie was primarily the CLIP-predictable component rather than the CLIP-residual component. Split-contribution analyses from M4 converged with this conclusion: PredCLIP exceeded ResCLIP in IntermediateVisual cortex and HighLevelVTC, but not EarlyVisual cortex (Supplementary Figure S3; Supplementary Tables S6-S7). M3 did not reliably improve prediction in EarlyVisual or IntermediateVisual cortex, but did improve prediction in HighLevelVTC (Δr_joint = .008 [.004, .012], q = .002.). In M4, including both calorie components simultaneously produced results that were numerically below M0 in IntermediateVisual cortex (Δr_joint = −.004, 95% CI [−.012, .004], q = 1.000) and not reliably above M0 in EarlyVisual cortex (Δr_joint = .001, 95% CI [−.003, .006], q = .493). This pattern likely reflects regularization instability when a noisy one-dimensional predictor (CalorieResCLIP) is fitted alongside a predictive one in a small stimulus set: the joint hyperparameter optimization may allocate weight suboptimally when one band carries negligible signal. Primary inference therefore rested on the nested M2 versus M0 and M2 versus M3 comparisons rather than on M4.

### Cortical flat maps visualized the spatial distribution of M0, M2, and M2-M0 gain

Surface maps were used to visualize the anatomical distribution of prediction accuracy and model gain (Figure 3). M0 showed strongest prediction in posterior visual cortex and intermediate occipitotemporal cortex (Figure 3B). M2 showed a similar broad spatial distribution but with increased prediction in ventral occipitotemporal cortex (Figure 3C). The M2-M0 difference map showed that the gain from CaloriePredCLIP was not confined to early visual cortex, but was most apparent in higher-level occipitotemporal regions overlapping with HighLevelVTC (Figure 3D). Statistical inference for the primary hypotheses was based on participant-level ROI analyses; the surface maps provide anatomical context for those effects.

**Figure 3.**
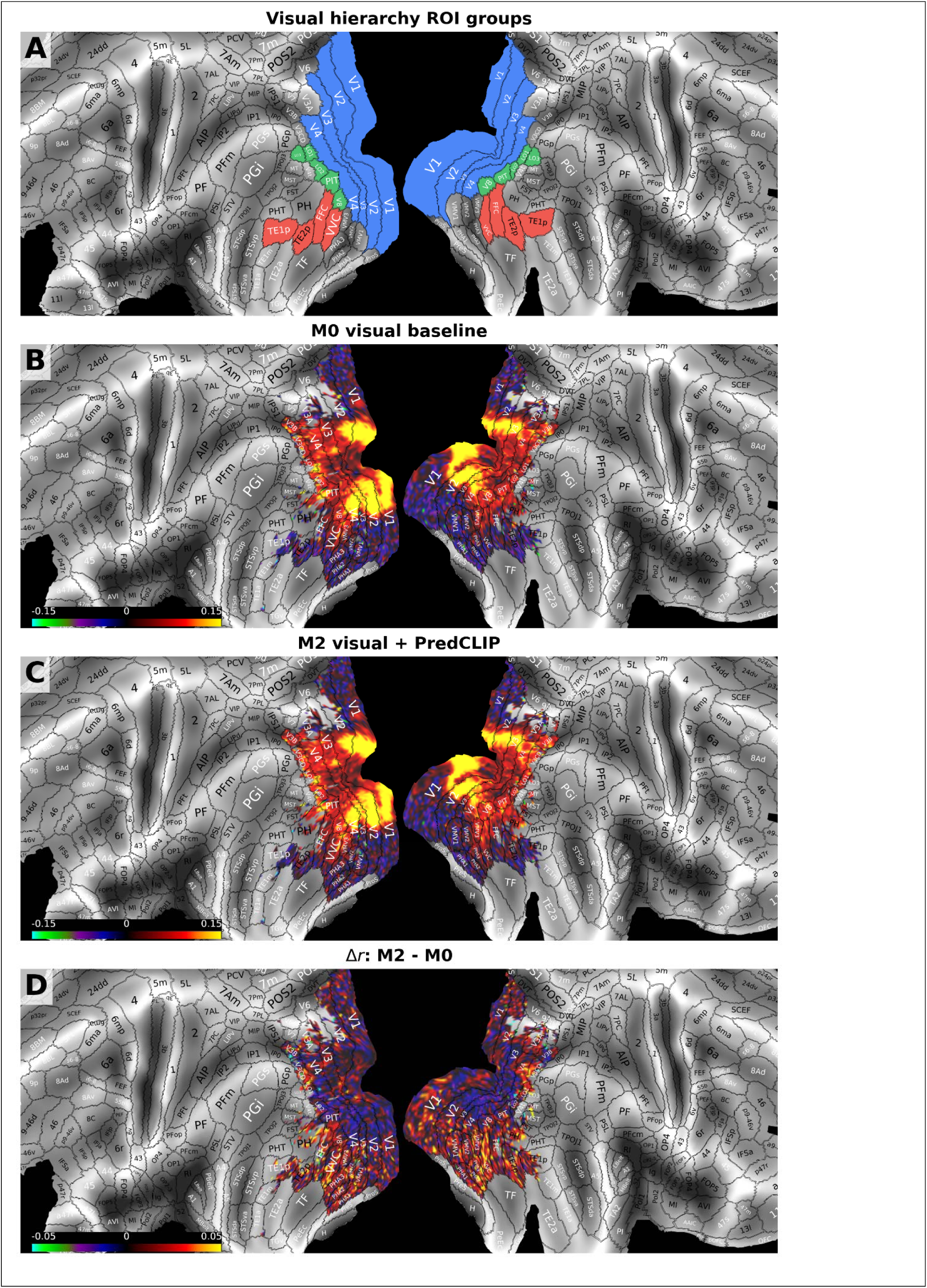
Surface distribution of M0, M2, and M2-M0 model gain. (A) Visual hierarchy ROI groups displayed on the HCP-MMP1.0 atlas surface. EarlyVisual is shown in blue, IntermediateVisual in green, and HighLevelVTC in red. (B) Group-average held-out prediction accuracy for M0. (C) Group-average held-out prediction accuracy for M2. (D) Difference map showing the gain from adding CaloriePredCLIP to M0, computed as M2 - M0. Surface maps visualize the spatial distribution of effects; primary statistical inference was performed at the participant level within the predefined ROIs and is reported in Figure 2 and Supplementary Tables S4-S5.

### Robustness and secondary analyses

Secondary and robustness analyses are reported in the supplementary materials. Full condition-shuffle controls produced null prediction values close to zero, indicating that the modeling procedure did not generate positive prediction when the stimulus-response mapping was destroyed (Supplementary Table S8). Calorie-label shuffle controls showed that the true M2 − M0 gain exceeded the shuffled-label mean in IntermediateVisual cortex and HighLevelVTC (Supplementary Table S8b), confirming that the CaloriePredCLIP gain in these regions depended on the true ordering of images along the calorie-aligned semantic axis. In EarlyVisual cortex, however, specificity was slightly negative (mean specificity = −.0003, 95% CI [−.0004, −.0001], Cohen’s dz = −.72) and did not support the predicted positive specificity effect under the directional test (q = .999). Thus, although M2 improved over M0 in EarlyVisual cortex, this gain should not be interpreted as evidence for a specific calorie-aligned signal. Instead, it is more consistent with the possibility that adding a low-dimensional CLIP-derived scalar can capture residual image structure in early visual cortex even when its ordering is not specifically aligned with perceived calorie. The evidence for a calorie-aligned semantic contribution is therefore strongest in IntermediateVisual cortex and HighLevelVTC.

We characterized CalorieResCLIP with a set of diagnostics (Supplementary Table S11-S14). CalorieResCLIP was highly reliable across independent rating subsamples (split-half Spearman ρ = .85; Spearman-Brown corrected = .92). It was not recoverable from any CLIP representation tested, including the original 50-dimensional ViT-B/32 ridge model, a nonlinear RBF readout of that space, the full 512-dimensional embedding, and the discarded higher-order CLIP components (all cross-validated R² ≤ 0). At the item level, CalorieResCLIP correlated positively with objective calorie category, perceived calorie, processedness, and preparation level, and negatively with perceived health, naturalness, and single-ingredient status (all |ρ| ≈ .3 to .4).

## Discussion

The present study asked why perceived calorie content predicts neural responses to food images. Using a nested family of banded ridge encoding models, we decomposed perceived calorie into a component predictable from CLIP image embeddings (CaloriePredCLIP) and a residual component not captured by this CLIP-based prediction (CalorieResCLIP), and tested their respective contributions to neural prediction beyond a visual-semantic baseline. The undecomposed perceived-calorie ratings (CalorieRaw) improved prediction of neural responses beyond this visual-semantic baseline. We then asked what carried this additional predictive information. Perceived calorie was strongly predictable from CLIP image embeddings, and its neural-predictive contribution was primarily carried by a visual-semantic axis separating raw or minimally prepared foods from processed, prepared, dessert-like, and energy-dense foods. The CalorieRaw improvement should therefore not be interpreted as evidence that visual cortex encodes caloric magnitude as an abstract nutritional variable. The primary neural-predictive component of perceived calorie was CaloriePredCLIP rather than CalorieResCLIP, although the latter contributed a smaller, reliable effect that we characterize below. Importantly, this does not imply that CaloriePredCLIP contains information absent from CLIP. Rather, it shows that a specific calorie-aligned direction within CLIP-readable food-image space predicts neural responses and becomes increasingly relevant along the ventral visual hierarchy. Below, we discuss how this finding constrains the interpretation of calorie-related neural effects, what it implies for the organization of food representations across the ventral visual hierarchy, and how the residual calorie component should be interpreted.

The central implication is that calorie-related prediction in visual cortex reflects a broader food-quality dimension rather than an abstract representation of caloric magnitude. In the present stimulus set, perceived-calorie and perceived-health ratings were nearly geometrically identical (RV = .907), indicating that the behavioral structure associated with perceived calorie was also strongly health-aligned. Accordingly, the CLIP-predictable component of perceived calorie (CaloriePredCLIP) should not be interpreted as calorie-specific. Instead, it captured a shared food-quality axis characterized by processing-and naturalness-related structure, distinguishing raw or minimally processed foods from prepared, processed, dessert-like, and energy-dense foods (Figure 1C–D). This axis was recoverable from CLIP image embeddings and was expressed most strongly in higher-level visual-semantic representations (Supplementary Figure S2).

This interpretation is consistent with evidence that processed and unprocessed foods elicit distinct neural responses even when caloric content is controlled(Coricelli, Toepel, Notter, Murray, & Rumiati, 2019), and with work suggesting that food concepts are organized partly by distinctions between natural/raw and transformed/prepared foods, supported by sensory and functional semantic properties (Rumiati & Foroni, 2016; Vignando et al., 2019). Convergent evidence comes from a searchlight RSA study(Avery et al., 2025) that identified a behavioral food dimension reflecting degree of processing and healthfulness (their first principal component, correlated r = .97 with processing ratings) represented in ventral occipitotemporal cortex, including fusiform, lateral occipital, and parahippocampal cortex. Notably, this relationship persisted after regressing out both perceived and actual fat content, indicating that the ventral-temporal food axis they identified reflects processedness and healthfulness rather than nutritional or caloric magnitude.

A closely related dimension emerges within our own data. In the companion RSA study using the same dataset(Marrazzo et al., 2026), the group-average axis along which perceived calorie and perceived health covary was reliably expressed across lateral and ventral occipitotemporal cortex (the lateral parcels LO1 to LO3 falling within the IntermediateVisual band, and ventral occipitotemporal cortex within the HighLevelVTC band of the present hierarchy), the same shared food-quality dimension that CaloriePredCLIP recovers here directly from image structure. Across three complementary approaches, behavioral similarity judgments (Avery et al., 2025), neural representational geometry (Marrazzo et al., 2026), and image-computable encoding (present study), the neurally relevant ventral visual food axis is therefore this shared processedness and health dimension rather than caloric magnitude per se.

The present encoding results complement these representational analyses by additionally showing that this shared axis is recoverable from image-computable visual-semantic structure (CLIP) and that its neural relevance increases along the ventral visual hierarchy. More broadly, it aligns with evidence that deep neural networks trained on large-scale visual or multimodal data recapitulate aspects of ventral visual representational geometry (Kietzmann et al., 2019; Marrazzo et al., 2026; Yamins & DiCarlo, 2016), and that high-level visual regions encode stimulus categories in semantically structured spaces reflecting both visual and conceptual similarity (Huth et al., 2012; Kriegeskorte et al., 2008).

This distinction is important because the present claim is not that CLIP merely provides an additional technical predictor of neural responses. The stronger conceptual point is that the across-image structure of perceived-calorie ratings is partly recoverable from high-level visual-semantic image information. In other words, perceived-calorie ratings do not isolate caloric magnitude: they covary with broader food-quality properties, including whether a food appears raw or transformed, natural or processed, and single-ingredient or prepared. CLIP provides a useful way to expose this structure because it captures semantic relations among images that are not captured by low-level visual statistics alone(Radford et al., 2021). CaloriePredCLIP should therefore be understood as a supervised, subjectively anchored projection of CLIP-readable image structure. Its value is explanatory: it identifies the part of perceived calorie that is available from the visual-semantic content of the image and tests whether that specific axis is neurally relevant.

The comparison with CalorieResCLIP reinforces this interpretation. If calorie-related neural prediction were primarily driven by residual perceived-calorie variance beyond the tested CLIP-readable image structure, one would expect CalorieResCLIP to carry stronger neural-predictive information. Instead, M2 outperformed M3 in both IntermediateVisual cortex (Δr = .014, q = .004) and HighLevelVTC (Δr = .008, q = .001). Thus, although CalorieResCLIP contributed a smaller reliable effect in HighLevelVTC, the dominant neural-predictive component of perceived calorie was the CLIP-aligned visual-semantic component rather than the residual component not captured by the tested CLIP-based prediction.

The gain associated with the CLIP-predictable food-quality component was not uniform across the visual hierarchy. The M2–M0 gain increased significantly from EarlyVisual to HighLevelVTC (slope = .007 Δr per step, q < .001), with a pairwise advantage of HighLevelVTC over both earlier regions. This hierarchical pattern is consistent with the known organization of the ventral visual stream, in which representations become progressively more abstract, category-selective, and tolerant to image variation along the posterior-to-anterior axis(DiCarlo et al., 2012; Grill-Spector & Weiner, 2014). The sensitivity of HighLevelVTC to this food-semantic axis is also consistent with recent evidence for food-selective responses in human ventral visual cortex (Jain et al., 2023; Khosla et al., 2022; Pennock et al., 2023), and with broader evidence that ventral temporal cortex organizes object representations along semantically meaningful dimensions such as animacy, real-world size, and category membership (Konkle & Caramazza, 2013).

The hierarchical pattern is important because it argues against a purely low-level visual explanation of the effect. If CaloriePredCLIP improved prediction merely because it captured simple image properties correlated with perceived calorie, such as color, contrast, or texture, the gain would be expected to be strongest in EarlyVisual cortex, where such features dominate neural responses. Instead, the gain was largest in HighLevelVTC, where visual representations are more strongly shaped by object identity, category structure, and semantic associations. This does not imply that early visual properties are irrelevant to food perception; rather, it suggests that the neural-relevant food-quality component captured by CaloriePredCLIP is most strongly expressed at a level of the visual hierarchy where food images are represented in terms of higher-order category structure and semantic meaning (Charest, Kievit, Schmitz, Deca, & Kriegeskorte, 2014; Connolly et al., 2012; Konkle & Caramazza, 2013; Kriegeskorte et al., 2008).

The weak performance of M0 in HighLevelVTC (r_joint = .001, q = .364) reflects how CLIP was expressed in the baseline rather than an insensitivity of this region to visual-semantic information. In M0, CLIP was pooled with AlexNet and CORnet-IT into a single PCA-compressed HighVis band. When instead CLIP was given its own band with independent regularization, baseline prediction in HighLevelVTC recovered from chance (D0, CLIP removed: r = 0.000, not significant) to reliably above zero (D1: r = 0.028; D1 > D0: Δr = 0.028, d_z = 2.11, q < .001; Supplementary Fig. S6, Supplementary Table S10). The failure of M0 in this region is therefore a consequence of joint feature compression, consistent with ventral temporal cortex containing distributed, overlapping object representations organized along multiple dimensions rather than a single compact code(Bracci & Op de Beeck, 2023; Bracci, Ritchie, & Op de Beeck, 2017; Haxby et al., 2001; Op de Beeck, Haushofer, & Kanwisher, 2008), within which a broad compressed baseline can fail to express a specific subjective relevant direction. Critically, CaloriePredCLIP improved prediction even over this fairly regularized CLIP band, and did so specifically in HighLevelVTC (D2 > D1: Δr = 0.008, d_z = 1.26, q < .001; null in EarlyVisual and IntermediateVisual). Because CaloriePredCLIP lies within CLIP’s representational span by construction, this gain indicates that HighLevelVTC preferentially weights the food-quality-related visual-semantic direction isolated by CaloriePredCLIP more strongly than regression over the full CLIP band allocates to it, rather than reflecting information beyond CLIP.

Although the CLIP-predictable food-quality component was the dominant source of calorie-related prediction, CalorieResCLIP also improved prediction beyond the visual baseline in high-level VTC (Δr = .008, q = .002). This effect was smaller than the gain observed for CaloriePredCLIP (M2 > M3, Δr = .008, q = .001), indicating that residual perceived-calorie variance was not the primary source of the effect. To better understand what this residual captured, we performed a series of diagnostic analyses (Supplementary Tables S11–S14). CalorieResCLIP showed high split-half reliability across independent rating subsamples (split-half Spearman ρ = .85; Spearman–Brown corrected = .92), indicating that it reflected stable stimulus-level structure rather than purely random rating noise. Conceptually, CalorieResCLIP captures the systematic discrepancy between perceived-calorie ratings and the values predicted from CLIP image embeddings. Its high reliability indicates that this discrepancy reflects stable stimulus-level structure, but does not determine whether it arises from food knowledge not adequately specified by the tested CLIP representation or from image-computable visual-semantic structure missed by the tested representations and readouts.

Across the tested CLIP feature spaces, CalorieResCLIP was not meaningfully recovered from the visual-semantic representations examined here, including the original 50-dimensional CLIP representation, a nonlinear RBF readout of the same space, the full 512-dimensional ViT-B/32 embedding, and the discarded higher-order CLIP principal components (all cross-validated R² ≤ 0; Supplementary Table S12). At the item level, CalorieResCLIP remained moderately associated with perceived calorie, objective calorie category, processedness, and preparation level, and negatively associated with perceived health and naturalness (Supplementary Table S13). Thus, the residual retained stable food-quality structure, but in a form that was not well captured by the tested CLIP-based readouts.

These diagnostics constrain, but do not resolve, the interpretation of CalorieResCLIP. One possibility is that it reflects stable food knowledge not well specified by visual appearance as represented by CLIP, such as beliefs about the energy density of specific foods that diverge from their apparent visual properties. Another possibility is that it reflects image-computable or semantic structure relevant to perceived calorie that the tested representations and readouts did not capture. Because non-recoverability was established only for the feature spaces and models examined here, CalorieResCLIP should not be interpreted as definitive evidence for appearance-independent nutritional knowledge.

For this reason, CalorieResCLIP is most appropriately interpreted as a secondary comparison model. Its predictive contribution indicates that some stable perceived-calorie variance outside the CLIP-predictable axis contributed to prediction in HighLevelVTC. However, this contribution was smaller than that of CaloriePredCLIP, and its interpretation remains less determinate. The principal result therefore remains that calorie-related prediction in ventral visual cortex was driven predominantly by the CLIP-aligned visual-semantic food-quality axis, rather than by residual perceived-calorie variance.

Several limitations constrain the interpretation. First, the stimulus set comprised only 96 images, limiting the statistical power of the multivariate baseline models and the generalizability of the calorie-axis characterization to the specific item sample used. Second, the sample was modest (n = 25) and restricted to healthy young women within a narrow BMI range, limiting generalizability across sexes and to broader populations with different demographic characteristics, dietary experience, or eating-related traits. In particular, it remains unclear to what extent the present representational profiles generalize to individuals with overweight or obesity. Although altered responses to food images have been reported in ventral-temporal and valuation/interoceptive regions, associations between BMI and neural responses or representations of food have not been observed consistently (Franssen, Jansen, van den Hurk, Roebroeck, & Roefs, 2020; Pimpini et al., 2022; Yang, Wu, & Morys, 2021). Because the decomposition target and the primary inferences are group-level, the modest sample bears most on the precision of the more complex model comparisons (in particular M4) rather than on the group-level effects themselves. Third, subjective ratings were collected after scanning in a single session, raising the possibility that the ratings do not fully reflect the representations active during fMRI; for the stable, stimulus-linked structure in ventral visual cortex this concern is comparatively mild. Relatedly, the rapid event-related design, with short presentation times and an orthogonal fixation task, may have favored stimulus-driven ventral-stream representations over slower, integrative signals. Fourth, the HighVis baseline combined AlexNet, CORnet, and CLIP into a single PCA-compressed feature band. This provided a compact visual-semantic baseline but may have limited the efficiency with which specific semantic directions within CLIP space were expressed. The present findings should therefore be interpreted as showing that a targeted CLIP-predictable calorie axis captured neural-relevant structure beyond the broad baseline used here, not as showing that this axis contains information absent from CLIP itself (the CLIP-separated diagnostic, Supplementary Figure S6, addresses this directly). Additionally, perceived calorie and perceived health were nearly identical in the present stimulus set (RV = .907; Supplementary Table S9), meaning that CaloriePredCLIP is empirically inseparable from a health-aligned CLIP projection on these 96 stimuli; the result should therefore be understood as evidence for a shared calorie, health, naturalness, and processedness semantic axis rather than for caloric magnitude specifically. Fifth, the reanalysis preserved the passive viewing paradigm of the original study, in which participants performed an orthogonal fixation task rather than evaluating foods explicitly. The extent to which the neural representations identified here reflect automatic, task-independent food processing versus incidental attention modulation cannot be fully resolved without a direct comparison between passive viewing and explicit evaluation.

The present findings clarify why perceived calorie content predicts neural responses to food images. Rather than supporting an abstract nutritional code in visual cortex, the results suggest that a substantial, neurally relevant part of this shared food-quality axis is recoverable from image-computable visual-semantic structure: perceived calorie is neurally predictive in large part because it is aligned with a dimension of food-image space that an image model can extract, here CLIP. This dimension reflected a shared food-quality axis co-varying with both perceived calorie and perceived health, separating foods according to broad differences in preparation, processing, naturalness, and energy-dense appearance. A smaller, reliable residual was not recoverable from the CLIP embeddings tested, and whether it reflects food knowledge unavailable from appearance or image-computable structure that a richer model would capture remains open. Its contribution increased along the ventral visual hierarchy, with the strongest effects in higher-level ventral temporal cortex. These findings situate calorie-related neural effects within the broader organization of visual-semantic representations, and suggest that subjective food ratings can index structured dimensions of image meaning rather than isolated nutritional variables. Future work should test whether similar dimensions contribute to subjective wanting and food choice, particularly under tasks that require explicit evaluation rather than passive visual attention.

## Materials and methods

### Dataset reuse and analytic rationale

The present study re-analysed the fMRI dataset reported in (Marrazzo et al., 2026). Participants, stimuli, subjective ratings, MRI acquisition, and basic preprocessing were identical to the original study and are summarized below only where relevant for the present analyses. The current study differed in its analytic approach: instead of comparing neural and model representational dissimilarity matrices, we re-estimated stimulus-specific responses using GLMsingle(Prince et al., 2022) on unsmoothed CIFTI grayordinate time series in fsLR surface space (Glasser et al., 2013) and fitted cross-validated banded-ridge encoding models (Dupré la Tour, Eickenberg, Nunez-Elizalde, & Gallant, 2022; Nunez-Elizalde et al., 2019).

### Participants

The original study included 33 female participants. After exclusions for incomplete scanning or acquisition inconsistencies, the original analysed dataset consisted of 25 participants who each completed two fMRI sessions. All participants provided written informed consent, and the original study was preregistered on AsPredicted and approved by the Ethics Committee of the Faculty of Psychology and Neuroscience of Maastricht University (ERCPN-252_67_04_2022). The present encoding-model reanalysis was not part of the original preregistration and should therefore be considered a secondary, hypothesis-driven analysis of the existing dataset. Full recruitment, exclusion, and preregistration details are reported in the supplementary materials.

**Table 1.** Participants’ characteristics. Reported are mean (M), standard deviation (SD), and range (minimum–maximum) for age, body mass index (BMI), time since last meal, and subjective hunger ratings. Hunger was assessed using a visual analogue scale (VAS; 0 = not hungry at all, 100 = very hungry). Time since last meal is reported in minutes.

| Variable | M | SD | Range (min – max) |
| --- | --- | --- | --- |
| Age | 21.28 | 2.74 | 18 – 31 |
| **BMI** | 21.74 | 1.47 | 19.60 – 24 |
| **Last eaten (min)** | 130.70 | 20.52 | 100 – 195 |
| **Hunger level**<br>**(VAS)** | 48.30 | 16.05 | 24 – 82 |

### Experimental design, stimuli, and data acquisition

Participants completed two fMRI sessions scheduled approximately one week apart and at a similar time of day. Before each scanning session, participants completed a hunger assessment to document metabolic state. In the second session, participants additionally completed an out-of-scanner stimulus-rating task and anthropometric measurements. Full procedural details are provided in the supplementary materials.

The stimulus set consisted of 96 food images selected from online sources and the Food-pics database (Blechert et al., 2019; Blechert et al., 2014). Images were placed on a uniform gray background (RGB: 191, 191, 191), subtended on average 3.7° × 2.8° of visual angle, and were presented centrally on the screen. The stimulus set was balanced to include 48 high-calorie and 48 low-calorie foods, with equal numbers of sweet and savory items within each calorie category. This yielded 24 high-calorie savory, 24 high-calorie sweet, 24 low-calorie savory, and 24 low-calorie sweet stimuli, forming a balanced stimulus space across calorie category and taste category. The trial structure and stimulus-space organization are shown in Figure 1A.

In the scanner, participants performed an orthogonal color-discrimination task adapted from a rapid event-related design (Kriegeskorte et al., 2008). On each trial, a food image was presented for 1.4 s with a blue or green fixation cross superimposed, and participants indicated the color of the fixation cross using an MR-compatible response box. The task was designed to maintain visual attention while avoiding explicit food evaluation during scanning. Each functional run contained all 96 food images, each presented once in randomized order, together with interspersed null events. Across seven runs per session and two sessions, each stimulus was presented 14 times in total. The intertrial interval was jittered and corresponded to 1, 2, or 3 repetition times (TR).

After scanning in the second session, participants rated all food stimuli on palatability, familiarity, perceived health value, and perceived caloric content using 100-mm visual analogue scales. These ratings provided the subjective variables used to construct the calorie-related predictors in the encoding analyses.

MRI data were acquired at Scannexus (Maastricht, The Netherlands) using a 3T Siemens MAGNETOM Prisma Fit scanner with a 64-channel head-neck coil. Functional images were acquired using a T2*-weighted multiband EPI sequence (TR = 1500 ms, TE = 30 ms, flip angle = 71°, field of view = 208 × 208 mm², voxel size = 2 mm isotropic, multiband factor = 3, GRAPPA factor = 2). Each run contained 410 volumes. High-resolution anatomical images were acquired using a T1-weighted MPRAGE sequence (TR = 2250 ms, TE = 2.21 ms, inversion time = 900 ms, flip angle = 9°, field of view = 256 × 256 mm², voxel size = 1 mm isotropic). In each session, one reversed phase-encoding spin-echo EPI image was acquired for susceptibility-distortion correction. Full acquisition and task details are reported in the supplementary materials.

### Preprocessing

Functional MRI data were preprocessed with fMRIPrep 25.2.2 ((Esteban et al., 2018; Esteban et al., 2019); RRID:SCR_016216), based on Nipype 1.10.0 ((Gorgolewski et al., 2011; Gorgolewski et al., 2018); RRID:SCR_002502). Preprocessing included correction for B0 field inhomogeneity using available reversed phase-encoding EPI images with *topup*(Andersson, Skare, & Ashburner, 2003), head-motion correction with MCFLIRT(Jenkinson, Bannister, Brady, & Smith, 2002), boundary-based registration of functional images to the T1-weighted anatomical reference(Greve & Fischl, 2009), anatomical segmentation and surface reconstruction with FreeSurfer(Dale, Fischl, & Sereno, 1999), and resampling to standard volumetric and surface spaces. The full fMRIPrep preprocessing description is reported in the supplementary materials.

For the present encoding analyses, we used the preprocessed CIFTI 91k grayordinate dtseries outputs in fsLR surface space (Glasser et al., 2013). Analyses were restricted to cortical surface grayordinates; subcortical grayordinates were not included. The CIFTI/fsLR representation was chosen because the encoding models were applied to cortical response patterns and because the HCP-MMP1.0 atlas used for region-of-interest definition is defined in fsLR surface space (Glasser et al., 2016). No additional spatial smoothing was applied before GLMsingle or encoding-model fitting, in order to preserve vertex-level response patterns. fMRIPrep-derived confound time series were generated and inspected, but no separate confound-regression step was applied before GLMsingle; denoising was performed within the GLMsingle framework described below.

### Stimulus-specific response estimation with GLMsingle

Stimulus-specific neural responses were estimated using GLMsingle (Prince et al., 2022), applied directly to the unsmoothed CIFTI 91k grayordinate time series. No additional spatial smoothing was applied, because the encoding analyses operated on distributed vertex-level response patterns.

For each run, the design matrix contained 96 stimulus regressors, one for each food image, with impulse events at stimulus onset. Null events were left as implicit baseline. All usable runs across the two scanning sessions were entered jointly, and a session-indicator vector was supplied to GLMsingle. GLMsingle was configured with HRF library fitting, GLMdenoise, fractional ridge regression, and percent BOLD signal change normalization enabled.

Trial-wise response estimates were extracted from the Type-D GLMsingle output, which combines HRF fitting, GLMdenoise, and fractional ridge regression. For most participants, this yielded 1344 trial-wise beta estimates, corresponding to 96 stimuli presented 14 times each. For two participants, one functional run was excluded because of missing stimulus trials, resulting in 13 usable runs and 1248 trial-wise beta estimates. These participants were retained in the main encoding analyses because each stimulus still had usable repeated estimates and complete model outputs. For each participant, trial-wise betas were averaged within stimulus identity to obtain one condition-level response pattern per image, yielding a 96 × vertices response matrix.

As a descriptive quality-control measure, we estimated repeated split-half noise ceilings from the repeated stimulus presentations. For each participant, the available repetitions of each stimulus were repeatedly divided into two approximately equal halves. Condition-level response estimates were computed separately for each half, and split-half reliability was estimated across the 96 stimulus conditions. Split-half correlations were averaged across repeated splits using Fisher-z transformation, Spearman–Brown corrected to estimate the reliability of the full condition-average response, and converted to an upper-bound prediction correlation by taking the square root of the non-negative corrected reliability. The resulting vertex-wise noise-ceiling estimates were averaged within each ROI for each participant; Figure 2 shows the group mean and 95% confidence interval across participants.

To ensure that all nested encoding models were evaluated in the same analysis space, a compact cortical vertex mask was constructed. Vertices were retained if they were present in at least 90% of participants and if their mean across-stimulus standard deviation exceeded the 50th percentile of the group distribution. This excluded vertices with poor coverage or minimal stimulus-related variability and yielded approximately 29,000–30,000 vertices per participant. Full GLMsingle implementation details are provided in the supplementary materials.

### Image-level feature spaces

For each of the 96 food images, feature representations were extracted and aligned to the canonical stimulus order used in the fMRI design. The primary model family used two multivariate visual feature bands, LowVis and HighVis, together with one-dimensional calorie-related predictors described below. LowVis was intended to capture low-level image-computable structure, whereas HighVis was intended to capture mid-to-high-level visual and semantic image structure.

LowVis combined Gabor energy features and color histogram features. Gabor features were included as a biologically motivated approximation of orientation- and spatial-frequency-selective responses in early visual cortex (Jones & Palmer, 1987; Kay et al., 2008; Nishimoto & Gallant, 2011). Color features were computed in CIELab space to capture chromatic structure in a perceptually interpretable color space. These features were grouped into a single LowVis band because both describe image structure available before object- or food-level semantic interpretation. Before concatenation, the Gabor and color feature matrices were kernel-normalized to unit trace; the concatenated matrix was then reduced to 80 principal components, retaining 99.4% of variance.

HighVis combined features from AlexNet (Krizhevsky, Sutskever, & Hinton, 2012), CORnet-S (Kubilius et al., 2019), and CLIP (Radford et al., 2021). AlexNet and CORnet-S were included as hierarchical object-vision models that capture intermediate and high-level visual representations relevant to the ventral visual stream (Yamins & DiCarlo, 2016). CLIP was included because contrastive image-text training yields embeddings that capture high-level visual-semantic structure beyond object-recognition features alone (Radford et al., 2021), and placing it in the baseline makes the calorie tests conservative: any calorie-related gain must exceed the visual-semantic structure CLIP already explains, rather than being credited with it. The HighVis components were grouped into a single visual-semantic baseline because the goal was not to adjudicate among deep-network architectures, but to construct a strong baseline against which calorie-related predictors had to explain additional variance. As for LowVis, all HighVis subcomponents were kernel-normalized before concatenation and PCA reduction to 80 components, retaining 94.9% of variance.

Full feature-extraction details, including layer selection, feature dimensionalities, normalization procedures, and feature-space diagnostics, are provided in the supplementary materials.

### Subjective calorie ratings and calorie decomposition

To define the subjective calorie predictor, we used participants’ perceived-calorie ratings collected after scanning. For each image, ratings were averaged across participants to obtain a single group-level perceived-calorie value, and this vector was z-scored across the 96 stimuli. This predictor was referred to as CalorieRaw, because it represents the original perceived-calorie rating vector before decomposition (see below). Group-mean rather than individual-participant ratings were used as the primary subjective target because the present analysis focused on the shared semantic component of food calorie perception that is consistent across participants, rather than on idiosyncratic individual differences in calorie knowledge or preference.

Perceived calorie should not be treated as a pure estimate of physical energy density: a neural effect of CalorieRaw could arise because the brain represents caloric content per se, or because calorie ratings index a broader image-computable dimension of food-image space correlated with perceived calorie.

To operationalize this distinction, we used CLIP image embeddings as a model of high-level visual-semantic image structure. Because CLIP is trained to align images with natural-language descriptions, its image embeddings provide a way to estimate the component of perceived calorie that is recoverable from image-computable semantic structure. The group-level perceived calorie ratings (CalorieRaw) were therefore decomposed into a CLIP-predictable component and a CLIP-residual component. CaloriePredCLIP was defined as the component of perceived calorie that could be predicted from CLIP image embeddings. This component captures the part of perceived calorie that is recoverable from the image’s visual-semantic structure as encoded by CLIP. CalorieResCLIP was defined as the residual perceived-calorie component after removing the CLIP-predictable part. This residual captures calorie-rating variance not recoverable from the CLIP embeddings used here, and therefore tests whether calorie-related neural prediction extends beyond what these embeddings capture. Because CLIP is a single vision-language model with a specific architecture, training set, and readout, the CLIP-recoverable component is a lower bound on image-computable structure, not its entirety. The residual therefore indexes variance not recoverable from the CLIP representations tested here, which may reflect either food knowledge unavailable from appearance or image-computable structure that a different or richer model would capture. By construction, CalorieRaw = CaloriePredCLIP + CalorieResCLIP.

For each participant, the decomposition target was a leave-one-participant-out group mean of perceived calorie ratings (the same quantity as CalorieRaw but excluding the participant being predicted), computed by averaging ratings across all other participants. This prevented the neural predictor for a given participant from being defined by that participant’s own subjective ratings, avoiding circularity between subjective and neural measurements.

Because encoding models were evaluated using 5-fold cross-validation across stimulus images (see *Encoding model fitting and evaluation* for details), the CLIP-based calorie decomposition was performed separately within each outer training fold. Within each fold, the calorie target was z-scored using training-image statistics only, and a regularized linear model was fitted to predict perceived calorie from CLIP features using only the training images. The fitted mapping was then applied to the held-out images to obtain fold-specific CaloriePredCLIP values; CalorieResCLIP was computed, within the z-scored target space, as the difference between the observed (z-scored) perceived-calorie target and the CLIP-predicted component.Thus, held-out images were never used to estimate the CLIP-to-calorie mapping or scaling parameters.

### Characterizing the calorie decomposition

#### CLIP-predicted component (CaloriePredCLIP)

To characterize what information was captured by CaloriePredCLIP, we performed complementary diagnostic analyses. First, we quantified how well different image-feature spaces predicted perceived calorie ratings. Perceived calorie was predicted from LowVis, HighVis, CLIP, and combined feature spaces using cross-validated ridge regression. Prediction accuracy was summarized using cross-validated explained variance, Pearson correlation, Spearman correlation, and the residual variance fraction. This analysis tested whether perceived calorie was more strongly recoverable from low-level image structure or from higher-level visual-semantic representations.

Second, we inspected the food images located at the low and high ends of the CaloriePredCLIP axis. This qualitative inspection was used to assess whether the axis corresponded to interpretable differences in food appearance, food category, or semantic structure. Because this step was descriptive, it was not used for statistical inference but served to guide interpretation of the learned CLIP-predictable calorie dimension.

Third, we related CaloriePredCLIP to CLIP text-probe axes capturing broad food-semantic contrasts. Text prompts were constructed to contrast concepts such as high-versus low-calorie foods, processed versus natural foods, prepared versus raw foods, and dessert-like versus produce-like foods. For each contrast, CLIP text embeddings were used to define a semantic axis, and food-image embeddings were projected onto that axis. These text-probe scores were then correlated with CaloriePredCLIP to determine which semantic contrasts best described the CLIP-predicted calorie component.

Finally, we examined whether processing/preparation-related structure was expressed more strongly in later CLIP representations than in earlier visual layers. This layerwise analysis tested whether the structure captured by CaloriePredCLIP reflected higher-level visual-semantic information rather than low-level image properties. Together, these analyses were used to characterize whether CaloriePredCLIP reflected an image-computable food-semantic axis rather than a pure estimate of nutritional calorie content.

#### CLIP-residual component (CalorieResCLIP)

To characterize the CLIP-residual calorie component, we performed a series of supplementary diagnostics. CalorieResCLIP was defined as the residual component of perceived calorie ratings after predicting group-average perceived calorie from CLIP image embeddings. First, we assessed whether this residual component reflected reliable subjective structure rather than rating noise. To this end, participants’ calorie ratings were split into two independent halves, and the full decomposition procedure was repeated separately in each half: calorie ratings were averaged within split, predicted from CLIP features using the same cross-validated ridge-regression procedure, and residualized to obtain two independent CalorieResCLIP estimates. The correlation between split-specific residual vectors was computed across stimuli and Spearman–Brown corrected to estimate full-sample reliability. We also expressed this reliability as a correlation-scale ceiling, defined as the square root of the positive Spearman–Brown-corrected reliability.

Second, we tested whether CalorieResCLIP could be recovered from richer or alternative CLIP representations. We predicted the final CalorieResCLIP vector using linear ridge regression from the original 50-dimensional ViT-B/32 ridge model, nonlinear kernel ridge regression from the same CLIP space, linear ridge regression from full 512-dimensional CLIP embeddings, and linear ridge regression from discarded higher-order CLIP components. Model performance was evaluated using cross-validated R², Pearson correlation, and Spearman correlation between predicted and observed CalorieResCLIP values.

Third, we examined the item-level structure of CalorieResCLIP by correlating it with subjective, categorical, manually annotated, and CLIP text-axis anchors. These anchors included perceived calorie, objective calorie category, perceived health, familiarity, processedness, naturalness, preparation level, raw-produce status, fruit/vegetable status, single-ingredient status, and contrastive CLIP text axes indexing processed versus natural foods, energy-dense treats versus light foods, and baked/confectionery foods versus fresh produce. Statistical inference for these item-level associations was obtained using item-label permutation of the CalorieResCLIP vector.

#### Encoding model family

Neural responses were predicted from image features using banded ridge regression(Nunez-Elizalde et al., 2019), implemented with the himalaya toolbox(Dupré la Tour et al., 2022). Banded ridge was used because the feature spaces differed substantially in dimensionality and structure. LowVis and HighVis were 80-dimensional multivariate spaces, whereas the calorie predictors were one-dimensional subjective or derived semantic axes. A standard ridge model with a single regularization parameter would regularize all features jointly and would therefore be inappropriate when feature spaces differ in dimensionality and collinearity. Banded ridge assigns each feature band its own regularization weight, allowing each band to contribute according to its predictive utility independently of its dimensionality.

Five nested models were fitted. M0 was the visual-semantic baseline and contained LowVis + HighVis. M1 added the undecomposed perceived-calorie ratings (CalorieRaw) to the visual baseline. M2 added the CLIP-predicted calorie component (CaloriePredCLIP) to the visual baseline and was the primary explanatory model. M3 added the CLIP-residual calorie component (CalorieResCLIP) and served as the specificity model. M4 included both decomposed calorie components and was used to estimate their relative split contributions in a common model. The model family is summarized in Table 2.

**Table 2.** Nested encoding-model family. The table summarizes the feature spaces included in each banded ridge encoding model and the primary question addressed by each model. All models were evaluated using the same cross-validation procedure and cortical analysis mask. M0 served as the visual-semantic baseline. M1 tested whether raw perceived calorie improved prediction beyond this baseline. M2 tested whether the CLIP-predictable component of perceived calorie improved prediction and was the primary explanatory model. M3 tested whether the CLIP-residual component carried additional predictive information. M4 included both decomposed calorie components and was used for split-contribution analyses.

| Model | Feature spaces | Primary question |
| --- | --- | --- |
| M0 | LowVis + HighVis | How much neural response structure is explained by the visual-semantic baseline? |
| M1 | LowVis + HighVis + CalorieRaw | Does group-level perceived calorie improve prediction beyond the visual-semantic baseline? |
| M2 | LowVis + HighVis + CaloriePredCLIP | Does the CLIP-predictable component of perceived calorie improve prediction beyond the baseline? |
| M3 | LowVis + HighVis + CalorieResCLIP | Does the component of perceived calorie not captured by CLIP improve prediction beyond the baseline? |
| M4 | LowVis + HighVis + CaloriePredCLIP + CalorieResCLIP | How do the CLIP-predictable and CLIP-residual components contribute when estimated in a common model? |

If CalorieRaw improves prediction beyond M0 but CaloriePredCLIP explains the same or most of this improvement, the calorie effect is likely carried by image-computable visual-semantic structure. If CalorieResCLIP improves prediction, this would indicate calorie-related variance beyond CLIP-accessible semantics. If CaloriePredCLIP outperforms CalorieResCLIP, the most parsimonious interpretation is that calorie-related neural prediction is driven primarily by an image-computable semantic dimension aligned with perceived calorie rather than by abstract caloric magnitude. Additionally, in the main model family, CLIP features were pooled with AlexNet and CORnet-IT and jointly reduced to 80 principal components within the HighVis band. To verify that the visual baseline (M0) was not artificially weakened by this joint compression, which could inflate the apparent gain of CaloriePredCLIP over M0, we fit a complementary diagnostic family in which CLIP was given its own band. D0 contained LowVis and a CLIP-free HighVis band (AlexNet + CORnet-IT only); D1 added CLIP as a separate band with its own regularization (the same 50-component ViT-B/32 representation from which CaloriePredCLIP was derived); D2 added CaloriePredCLIP on top of D1. Because D0 ⊂ D1 ⊂ D2, D1 − D0 isolates CLIP’s unique contribution and D2 − D1 tests whether the calorie-aligned axis improves prediction beyond a fairly regularized CLIP band. CaloriePredCLIP lies within the span of the CLIP band by construction, so a positive D2 − D1 reflects preferential weighting of the calorie-aligned direction rather than information outside CLIP. Fitting, nested cross-validation, per-band regularization, and ROIs were identical to the main analysis.

Palatability and familiarity were not included in the final nested model family. In preliminary model versions, palatability was included as an additional subjective band after residualizing it with respect to familiarity and perceived calorie, and familiarity was included as a nuisance band. These analyses were useful for checking whether the calorie-related effects could be attributed to general food liking or stimulus familiarity. However, these predictors were not central to the present hypothesis and did not alter the main calorie-decomposition pattern. To keep the final model family targeted and interpretable, we therefore restricted the primary analyses to the predictors needed to adjudicate the central question: whether the neural-relevant component of perceived calorie is the CLIP-predictable component or the CLIP-residual component.

#### Encoding model fitting and evaluation

Model performance was evaluated using 5-fold outer cross-validation across the 96 stimulus conditions. Each fold used approximately 77 images for training and 19 images for testing. Cross-validation was performed across stimulus identities, so that prediction accuracy reflected generalization to held-out food images rather than prediction of repeated measurements of the same image. Five folds were chosen as a compromise between maximizing the number of training images available for model fitting and retaining enough held-out images per fold to estimate prediction accuracy reliably.

Within each outer training fold, all feature preprocessing and model fitting steps were performed using training images only. Feature matrices were standardized to zero mean and unit variance per feature dimension, and the resulting scaling parameters were applied to the held-out images. For each feature band, a linear kernel was computed as *K* =*XX^T^* Kernels were then double-centered and Frobenius-normalized, placing all feature bands on a common scale before banded-ridge fitting. This ensured that band weights reflected predictive utility rather than raw feature variance, dimensionality, or kernel magnitude.

For each outer fold, banded-ridge hyperparameters were optimized within the training set using an inner cross-validation procedure in two stages. First, 100 random-search initializations were evaluated using 5-fold inner cross-validation on the training images. Second, the best initialization was refined using hypergradient descent. This random-search-plus-hypergradient strategy was used because random search alone may not identify the optimal banded-ridge parameter allocation precisely, whereas hypergradient descent alone can be sensitive to initialization (Dupré la Tour et al., 2022).

Held-out prediction accuracy was quantified at each vertex as the Pearson correlation between the predicted and observed condition-averaged beta responses for held-out images, computed within each outer fold and then averaged across folds. This cross-validated whole-model prediction accuracy is referred to as r_joint, where “joint” indicates that the prediction is generated by the full set of feature bands included in a given model. For example, r_joint for M0 reflects the joint prediction from LowVis and HighVis, whereas r_joint for M2 reflects the joint prediction from LowVis, HighVis, and CaloriePredCLIP.

For M4, per-band split contributions were computed to estimate how LowVis, HighVis, CaloriePredCLIP, and CalorieResCLIP contributed when included in a common model. These split scores were treated as secondary descriptive measures; primary inference was based on nested model gains rather than split scores.

#### Region-of-interest definition and hierarchy analysis

Regions of interest were defined using the HCP-MMP1.0 atlas in fsLR-91k surface space (Glasser et al., 2016). This atlas was chosen because it provides anatomically and functionally defined cortical parcels in the same surface space as the CIFTI data, allowing ROI assignment without volumetric-to-surface resampling.

The primary analyses focused on three ROI groups spanning a posterior-to-anterior visual hierarchy, motivated by evidence that visual representations become progressively more complex, category-relevant, and semantically structured along the ventral stream (DiCarlo et al., 2012; Grill-Spector & Weiner, 2014). “EarlyVisual” included V1, V2, V3, and V4, covering primary and early extrastriate visual cortex expected to be sensitive to low-level image properties such as contrast, orientation, spatial frequency, color, and local texture (Wang, Mruczek, Arcaro, & Kastner, 2015). “IntermediateVisual” included V8, PIT, LO1, LO2, and LO3, covering intermediate occipitotemporal regions implicated in higher-order visual analysis, object structure, and category-level processing (Larsson & Heeger, 2006; Wang et al., 2015). “HighLevelVTC” included FFC, VVC, TE1p, and TE2p, covering higher-level ventral temporal cortex where responses are expected to be increasingly organized by object- and category-level structure rather than by local image properties alone (Grill-Spector & Weiner, 2014). The three ROI groups are visualized on a cortical flat map in Figure 3A.

For hierarchy analyses, the ROI groups were assigned ordinal levels: EarlyVisual = 0, IntermediateVisual = 1, and HighLevelVTC = 2. This ordering was used to test whether model gains increased along the visual hierarchy. For each participant and ROI, prediction accuracy was averaged across vertices that were included in the compact cortical analysis mask and had finite model values. For each participant, model gain was computed within each ROI as the difference in held-out prediction accuracy between two nested models, for example M2 − M0. A linear slope was then fitted across the three ROI levels for each participant, and the resulting participant-level slopes were tested at the group level. A positive slope indicated that model gain increased from early visual cortex to higher-level ventral temporal cortex.

#### Statistical inference

All group-level inference used the participant as the inferential unit. For each ROI and model, participant-level r_joint values were averaged across valid vertices within the ROI and Fisher-z transformed before statistical testing. For paired model comparisons, the tested quantity was the within-participant Fisher-z difference between two models. Fisher-z transformation was applied because Pearson correlations are bounded on the raw r scale and have a non-normal sampling distribution, especially when values deviate from zero.

Group-level inference was performed using sign-flipping permutation tests. The sign-flipping test assumes that, under the null hypothesis of no group-level effect, participant-level values are symmetrically distributed around zero and are therefore exchangeable with respect to sign. For one-sample tests of r_joint > 0, this means that ROI-level prediction accuracies would be equally likely to be positive or negative under the null hypothesis of no reliable stimulus-response correspondence. For paired model comparisons, the sign-flip test was applied to within-participant Fisher-z differences, testing whether the gain of one model over another was consistently greater than zero across participants.

Monte Carlo sign-flipping was performed with 50,000 permutations, following standard non-parametric permutation-inference procedures for neuroimaging data (Nichols & Holmes, 2002; Winkler, Ridgway, Webster, Smith, & Nichols, 2014). One-sided p-values were computed using the Phipson-Smyth continuity correction, *p* =(exceed+1)/(M+1), where is the number of permutations (Phipson & Smyth, 2010). One-sample tests assessed whether each model produced above-zero prediction within each ROI. Paired comparisons tested the pre-specified nested model contrasts, including the primary contrast M2 > M0 and the key specificity contrast M2 > M3. Hierarchy analyses tested whether model gains increased across EarlyVisual, IntermediateVisual, and HighLevelVTC using participant-level gain slopes.

Multiple comparisons were controlled using the Benjamini-Hochberg false discovery rate procedure at .05. FDR correction was applied separately within predefined test families: one-sample model-performance tests, paired model-comparison tests, split-contribution tests, and hierarchy pairwise contrasts.

#### Permutation controls

Two permutation controls were performed. First, a full condition-shuffle control tested whether the cross-validation and regularization procedure produced positive prediction when the stimulus-response correspondence was destroyed. For each participant, the mapping between condition labels and neural response patterns was randomly permuted before model fitting, thereby preserving the structure of the fitting procedure while breaking the relationship between image features and neural responses.

Second, a calorie-label shuffle tested whether the M2 − M0 gain depended on the true alignment between food images and CaloriePredCLIP values, rather than on the generic benefit of adding a low-dimensional scalar predictor. In this control, visual features and neural responses remained aligned, but CaloriePredCLIP values were randomly permuted across images. For each participant, specificity was defined as the true M2 − M0 gain minus the mean shuffled M2 − M0 gain. Group-level inference on this specificity index was performed using sign-flipping across participants.

Together, these controls tested whether the encoding procedure generated positive prediction under destroyed stimulus-response correspondence, and whether the CaloriePredCLIP gain depended on the specific alignment between food images and their CaloriePredCLIP values rather than simply reflecting the addition of a one-dimensional CLIP-derived predictor with arbitrary image labels. Full implementation details are reported in the supplementary materials.

## Data availability

The raw neuroimaging and behavioural data used in this study are available through the DataverseNL repository associated with the companion study [https://doi.org/10.34894/TVPLVR]. Data and derived outputs specific to the present analyses are being prepared for public release and will be made available upon publication.

## Code availability

Code specific to the present analyses is being prepared for public release and will be made available upon publication.

## Acknowledgements

We thank Prof. Dr. Federico De Martino and Dr. Giancarlo Valente for helpful discussions on earlier versions of the manuscript. This study was financed by starter grant of the Dutch Ministry of Education, Culture and Science (OCW) (SG2024-5) awarded to Dr. Leonardo Pimpini and by a VIDI grant of the Dutch Research Council (NWO) (016.165.356) awarded to Prof. Dr. Anne Roefs.

## CRediT authorship contribution statement

**Giuseppe Marrazzo**: Conceptualization, Methodology, Data Curation, Validation, Formal Analysis, Visualization, Software, Writing – original draft, Writing – review & editing. **Anne Roefs**: Funding acquisition, Conceptualization, Writing – review & editing. **Leonardo Pimpini**: Project administration, Conceptualization, Methodology, Investigation, Funding acquisition, Writing – original draft, Writing – review & editing.

