## Supplementary Material for "Vision-language encoding models reveal an image-computable food-quality dimension in human occipitotemporal cortex"

### Supplementary Materials

### Materials and Methods

#### Participants

Thirty-three female participants volunteered for the original fMRI study. Only female participants were recruited to reduce variability associated with reported sex differences in neural responses to visual food cues. Participants were recruited through paper flyers, social media, and the Maastricht University participant recruitment system (SONA). One participant was excluded because she experienced sickness or claustrophobia during MRI scanning and did not complete the experiment.

Seven additional participants were excluded because of an acquisition inconsistency. These participants were scanned with a left–right phase-encoding direction, whereas the remaining participants were scanned with an anterior–posterior phase-encoding direction. This difference produced substantially different susceptibility-induced distortion patterns that could not be reliably corrected or harmonized across participants. These participants were therefore excluded to ensure consistency of preprocessing, spatial normalization, and multivariate pattern analyses.

The original analysed dataset consisted of 25 participants, each of whom completed two fMRI sessions. The original study was preregistered on AsPredicted and approved by the Ethics Committee of the Faculty of Psychology and Neuroscience of Maastricht University (ERCPN-252_67_04_2022). Before participation, all participants provided written informed consent and were compensated with either 5 university credits or a 40 EUR voucher.

For the present encoding-model reanalysis, one additional participant was excluded because fMRIPrep preprocessing did not produce usable CIFTI fsLR grayordinate outputs. The final inferential sample therefore consisted of 25 participants with complete GLMsingle outputs and complete model outputs across the full nested model family. The present encoding-model reanalysis was not part of the original preregistration and should therefore be considered a secondary, hypothesis-driven analysis of the existing dataset.

#### Experimental procedure

Participants completed two experimental sessions scheduled approximately one week apart and at a similar time of day. Approximately two weeks before the first scanning session, participants underwent a general screening for eligibility and MRI compatibility. At the start of each session, participants received scanner-safety instructions and task instructions and completed a hunger assessment to document metabolic state. In the second session, participants additionally completed a stimulus-rating task of approximately 25 minutes after scanning, and their height and weight were measured to calculate body mass index (BMI). Each session required approximately two hours in total. Participants received compensation and a debriefing letter at the conclusion of the study.

**Anthropometric and behavioral measurements**

Height, weight, and age were measured to compute BMI (kg/m$^2$). Participants were asked to remove their shoes and to wear jeans or trousers and a T-shirt during measurement. The same scale and measuring tape were used throughout data collection.

To standardize metabolic state across participants, individuals were instructed to eat a small snack, such as a sandwich and a piece of fruit, exactly two hours before each session and to refrain from eating or drinking, except water, until the session ended. Upon arrival, participants completed a three-item paper hunger questionnaire recording: (1) the time elapsed since the last eating moment, in minutes; (2) a brief description of the last food consumed; and (3) subjective hunger, rated on a 100-mm visual analogue scale ranging from 0 = not hungry at all to 100 = very hungry.

After scanning in the second session, participants rated each food stimulus along four dimensions: palatability (0 = not tasty at all, 100 = very tasty), familiarity (0 = completely unfamiliar, 100 = very familiar), perceived health value (0 = very unhealthy, 100 = very healthy), and perceived caloric content (0 = very low in calories, 100 = very high in calories). Ratings were collected in a blocked design, such that all stimuli were rated on one dimension within a block. Each block was followed by a short break, after which participants proceeded to the next rating dimension. The order of rating blocks was randomized across participants. Within each block, food stimuli were presented individually in randomized order and rated using a 100-mm visual analogue scale. This procedure ensured that each stimulus received one rating per dimension from each participant.

**Stimuli**

The stimulus set consisted of 96 food images selected from online sources, including Google and iStockphoto, and from the Food-pics database issued by the University of Salzburg (Blechert, Meule, Busch, & Ohla, 2014; Blechert, Lender, Polk, Busch, & Ohla, 2019). Images were formatted using Adobe Photoshop CS5.1 to obtain a standard resolution of 96 pixels/inch, and MATLAB was used to place each stimulus on a uniform gray background (RGB: 191, 191, 191). Stimuli subtended, on average, 3.7° × 2.8° of visual angle and were presented centrally on the screen.

The stimulus set was deliberately balanced to include 48 high-calorie and 48 low-calorie food items. Within each calorie category, the set contained equal numbers of sweet and savory foods, yielding 24 high-calorie savory, 24 high-calorie sweet, 24 low-calorie savory, and 24 low-calorie sweet stimuli. This balanced structure was relevant for interpreting category-level controls and the calorie-label permutation control reported in the main manuscript.

**MRI acquisition**

Brain imaging was performed at Scannexus (Maastricht, The Netherlands) using a 3T Siemens MAGNETOM Prisma Fit MRI scanner (Siemens Healthineers, Erlangen, Germany) equipped with a 64-channel head-neck coil. Participants lay supine in the scanner with their head stabilized using foam padding to minimize motion. Visual stimuli were presented via a mirror mounted on the head coil.

High-resolution anatomical images were acquired using a three-dimensional T1-weighted magnetization-prepared rapid gradient-echo sequence (MPRAGE; TR = 2250 ms, TE = 2.21 ms, inversion time = 900 ms, flip angle = 9°, field of view = 256 × 256 $mm^2$, voxel size = 1 × 1 × 1 $mm^3$). Functional images were acquired using a T2*-weighted multiband gradient echo-planar imaging sequence (TR = 1500 ms, TE = 30 ms, flip angle = 71°, field of view = 208 × 208 $mm^2$, voxel size = 2 × 2 × 2 $mm^3$). Whole-brain coverage was obtained with contiguous axial slices using multiband acceleration factor 3 and in-plane GRAPPA acceleration factor 2. Slices were acquired in an interleaved order. For each functional run, 410 volumes were collected.

Each participant completed two scanning sessions. In each session, one spin-echo EPI image with reversed phase-encoding direction was acquired and used to estimate susceptibility-induced distortions for all functional runs within that session. Two T1-weighted images acquired across sessions were combined to form a single high signal-to-noise anatomical reference.

### fMRI task

During functional imaging, participants performed a color-discrimination task adapted from a rapid event-related fMRI paradigm (Kriegeskorte et al., 2008). Each functional run lasted approximately 11 minutes and comprised a sequence of 96 food images. Each image was displayed for 1.4 s, and each stimulus appeared once per run. Across seven runs per session and two sessions in total, each stimulus was presented 14 times. Within each run, the order of stimulus presentation was fully randomized.

Participants were instructed to indicate the color of a fixation cross, blue or green, superimposed on each food image by pressing a button on an MR-compatible response box. This task was orthogonal to the food properties of interest and was intended to maintain visual attention without requiring explicit food evaluation during scanning. To avoid systematic response-mapping biases, fixation-cross color was pseudo-randomized across runs such that each food stimulus was paired equally often with each color across the experiment, seven times with a blue cross and seven times with a green cross. Button-color mappings were counterbalanced across participants.

Each functional run included null events to establish an implicit baseline. Four null events were presented at the beginning of each run, four at the end, and 32 were randomly interspersed among the food stimuli. Null events consisted of a centrally presented black fixation cross on a gray background. The intertrial interval was jittered and corresponded to 1, 2, or 3 repetition times. Stimulus presentation and response collection were controlled using Presentation software (Neurobehavioral Systems, Albany, CA, USA).

### Preprocessing

Functional MRI data were preprocessed using fMRIPrep 25.2.2 (Esteban et al., 2018, 2019; RRID:SCR_016216), which is based on Nipype 1.10.0 (Gorgolewski et al., 2011, 2018; RRID:SCR_002502). The following section provides the full preprocessing description for the dataset used in the present reanalysis. The main encoding analyses used the fMRIPrep-generated CIFTI 91k grayordinate dtseries outputs in fsLR surface space. Analyses were restricted to cortical grayordinates; subcortical grayordinates were not included. No additional spatial smoothing was applied before GLMsingle estimation or encoding-model fitting. fMRIPrep-derived confound regressors were generated and inspected, but no separate confound-regression step was applied before GLMsingle. Instead, denoising was performed within GLMsingle using its GLMdenoise procedure.

#### *Preprocessing of B0 inhomogeneity mappings*

A total of 14 fieldmaps were available within the input BIDS structure. A B0 non-uniformity map, or fieldmap, was estimated based on two or more echo-planar imaging references with reversed phase-encoding directions using *topup* (Andersson, Skare, & Ashburner, 2003; FSL).

*Anatomical preprocessing*

Two T1-weighted images were corrected for intensity non-uniformity using N4BiasFieldCorrection (Tustison et al., 2010), distributed with ANTs 2.6.2 (Avants et al., 2008; RRID:SCR_004757), and were used as the T1w anatomical reference throughout the workflow. The T1w reference was skull-stripped using a Nipype implementation of the antsBrainExtraction.sh workflow from ANTs, using OASIS30ANTs as the target template. Brain tissue segmentation of cerebrospinal fluid, white matter, and gray matter was performed on the brain-extracted T1w image using FAST (FSL; Zhang, Brady, & Smith, 2001).

An anatomical T1w reference was computed after registration of the two intensity-corrected T1w images using mri_robust_template (FreeSurfer 7.3.2; Reuter, Rosas, & Fischl, 2010). Brain surfaces were reconstructed using recon-all (FreeSurfer 7.3.2; Dale, Fischl, & Sereno, 1999; RRID:SCR_001847), and the brain mask estimated previously was refined using a custom variation of the Mindboggle method to reconcile ANTs-derived and FreeSurfer-derived cortical gray-matter segmentations (Klein et al., 2017; RRID:SCR_002438).

Volume-based spatial normalization to MNI152NLin6Asym and MNI152NLin2009cAsym standard spaces was performed through nonlinear registration with antsRegistration (ANTs 2.6.2), using brain-extracted versions of both the T1w reference and the T1w template. Templates were accessed with TemplateFlow 25.0.4 (Ciric et al., 2022). The selected templates were FSL's MNI ICBM 152 non-linear 6th Generation Asymmetric Average Brain Stereotaxic Registration Model (Evans et al., 2012; TemplateFlow ID: MNI152NLin6Asym) and the ICBM 152 Nonlinear Asymmetrical template version 2009c (Fonov et al., 2009; TemplateFlow ID: MNI152NLin2009cAsym).

*Functional preprocessing*

For each of the 14 BOLD runs per subject, a reference volume was generated by fMRIPrep for use in head-motion correction. Head-motion parameters with respect to the BOLD reference, including transformation matrices and six corresponding rotation and translation parameters, were estimated before any spatiotemporal filtering using

MCFLIRT (FSL; Jenkinson et al., 2002). The estimated fieldmap was aligned with rigid registration to the target EPI reference run, and the field coefficients were mapped onto the reference EPI using the resulting transform.

The BOLD reference was co-registered to the T1w reference using bbregister (FreeSurfer), which implements boundary-based registration (Greve & Fischl, 2009). Co-registration was configured with six degrees of freedom.

Several confounding time series were calculated from the preprocessed BOLD data, including framewise displacement, DVARS, and three region-wise global signals extracted from cerebrospinal fluid, white matter, and whole-brain masks. Framewise displacement was computed using both the Power formulation, defined as the absolute sum of relative motions (Power et al., 2014), and the Jenkinson formulation, defined as the relative root mean square displacement between affines (Jenkinson et al., 2002). DVARS was calculated according to the definitions implemented in Nipype following Power et al. (2014).

Physiological nuisance regressors were extracted using component-based noise correction (CompCor; Behzadi et al., 2007). Principal components were estimated after high-pass filtering the preprocessed BOLD time series using a discrete cosine filter with a 128-s cutoff. Components were estimated for both temporal CompCor and anatomical CompCor. Temporal CompCor components were calculated from the top 2% most variable voxels within the brain mask. For anatomical CompCor, probabilistic masks for cerebrospinal fluid, white matter, and combined cerebrospinal fluid plus white matter were generated in anatomical space. Instead of eroding these masks by two pixels in BOLD space, as in the original implementation, fMRIPrep subtracted a mask of voxels likely to contain gray-matter volume fraction. This mask was obtained by dilating a gray-matter mask extracted from the FreeSurfer aseg segmentation, ensuring that components were not extracted from voxels containing a minimal fraction of gray matter. The masks were resampled into BOLD space and binarized at 0.99. Components were also calculated separately within the white matter and cerebrospinal fluid masks. For each CompCor decomposition, the components with the largest singular values were retained such that the retained components explained 50% of the variance across the corresponding nuisance mask; remaining components were discarded.

The head-motion estimates were included in the corresponding fMRIPrep confounds file. Confound time series derived from head-motion estimates and global signals were expanded with temporal derivatives and quadratic terms (Satterthwaite et al., 2013). Frames exceeding 0.5 mm framewise displacement or 1.5 standardized DVARS were annotated as motion outliers. Additional nuisance time series were calculated using principal component analysis of the signal within a thin crown of voxels around the edge of the brain, as proposed by Patriat, Reynolds, and Birn (2017).

The BOLD time series were resampled onto the fsnative and fsaverage surfaces and onto the left-right symmetric fsLR template using Connectome Workbench (Glasser et al., 2013). CIFTI grayordinate dtseries files containing 91k samples were generated, with cortical surface data transformed directly to fsLR space and subcortical data transformed to 2-mm resolution MNI152NLin6Asym space. All resampling was performed in a single interpolation step by composing the relevant transformations, including head-motion transformations, susceptibility distortion correction, and anatomical co-registration transformations. Gridded volumetric resampling was performed using nitransforms with cubic B-spline interpolation. Non-gridded surface resampling was performed using mri_vol2surf (FreeSurfer). Many internal operations of fMRIPrep used Nilearn 0.10.2 (Abraham et al., 2014; RRID:SCR_001362).

For the present encoding-model reanalysis, only the cortical portion of the CIFTI 91k grayordinate dtseries outputs was used. This yielded 59,412 cortical grayordinates before subsequent GLMsingle-derived masking. Subcortical grayordinates were excluded because the primary hypotheses concerned cortical visual representations along the ventral visual hierarchy, and because the region-of-interest analyses used the HCP-MMP1.0 cortical atlas in fsLR surface space (Glasser et al., 2016).

**image-level feature extraction and feature-space construction**

For each of the 96 food images, multiple image-level feature representations were extracted and aligned to the canonical stimulus order used in the fMRI design. Feature spaces were designed to span low-level image-computable structure, higher-level visual and semantic structure, and food-relevant behavioral dimensions. The primary encoding-model family used two multivariate visual feature bands, LowVis and HighVis, together with one-dimensional calorie-related predictors described in the main Methods.

All feature matrices were constructed with one row per food image. Before model fitting, all feature matrices were aligned to the same 96-image order used to construct the GLMsingle design matrices and condition-level beta estimates.

*Low-level visual feature space: LowVis*

The LowVis feature space was designed to capture image-computable properties available before object- or food-level semantic interpretation. It combined Gabor energy features and color histogram features.

*Gabor features*

Gabor features were included as a biologically motivated approximation of orientation- and spatial-frequency-selective responses in early visual cortex (Jones & Palmer, 1987; Kay et al., 2008). Each food image was converted to grayscale and resized to 256 × 256 pixels. A bank of two-dimensional Gabor filters was applied across eight orientations and three spatial frequencies. Phase-invariant Gabor energy maps were computed and spatially pooled within a 4 × 4 grid, yielding a 384-dimensional feature vector for each image.

The resulting Gabor feature matrix had shape 96 × 384, with rows corresponding to images and columns corresponding to pooled orientation/spatial-frequency energy features.

*Color features*

Color features were computed to capture chromatic structure that may be relevant for food perception and may covary with perceived caloric content. Images were represented in CIELab color space rather than RGB because CIELab is approximately perceptually uniform and separates luminance from chromatic dimensions. For each image, a three-dimensional color histogram was computed using eight bins for each Lab channel, yielding 8 × 8 × 8 = 512 histogram bins. Histograms were normalized to unit sum, resulting in a 512-dimensional color feature vector for each image.

The resulting color feature matrix had shape 96 × 512.

*LowVis band construction*

Gabor and color features were grouped into a single LowVis band because both describe pre-semantic image structure. Treating them as separate bands would risk attributing shared low-level image variance to one subcomponent artifactually. Before concatenation, each feature block was centered and kernel-normalized to unit trace. Specifically, for each feature matrix $X$, the linear kernel $K = XX^{\top}$ was computed after centering, and the feature matrix was rescaled such that the trace of the kernel was equal to one. This ensured that no subcomponent dominated the concatenated LowVis representation by virtue of raw variance magnitude rather than information content.

The kernel-normalized Gabor and color matrices were concatenated and reduced using principal component analysis. The resulting LowVis band contained 80 principal components and retained 99.4% of the variance of the concatenated low-level feature space. The final LowVis matrix had shape 96 × 80.

*High-level visual-semantic feature space: HighVis*

The HighVis feature space was designed to capture mid-to-high-level visual and semantic image structure. It combined features from AlexNet, CORnet-S, and CLIP. These models were included because they capture complementary levels of visual representation: hierarchical object-recognition features, biologically constrained ventral-stream-like features, and language-aligned visual-semantic embeddings.

*AlexNet features*

AlexNet features were extracted from a pretrained AlexNet model (Krizhevsky et al., 2012). Mid-level visual features were extracted from convolutional layers 3 and 5, which capture intermediate object-vision representations. High-level features were extracted from the fully connected layer fc6, which captures more global object-level structure. These layers were selected because hierarchical convolutional network layers have been widely used as computational models of progressively more complex visual representations along the ventral visual stream (Yamins & DiCarlo, 2016).

The AlexNet mid-level feature representation consisted of conv3 and conv5 features reduced to 50 principal components. The AlexNet high-level representation consisted of fc6 features reduced to 50 principal components.

*CORnet-S features*

CORnet-S features were included as a biologically constrained model of the primate ventral visual stream (Kubilius et al., 2019). The present analyses used IT-stage activations, intended to approximate high-level inferotemporal representations. These features were used because the study focused on visual cortical representations that may become increasingly semantic and category-relevant along the ventral hierarchy.

The CORnet-S IT feature matrix was aligned to the same 96-image stimulus order and entered as a high-level visual feature block.

*CLIP features*

CLIP features were extracted using a pretrained CLIP ViT-B/32 image encoder (Radford et al., 2021). Each food image was passed through the CLIP image encoder to obtain a 512-dimensional image embedding. CLIP was included because contrastive image–text training yields visual embeddings that capture semantic and categorical structure beyond standard object-recognition features. These embeddings are therefore well suited for characterizing the high-level visual-semantic structure of food images.

The 512-dimensional CLIP image embeddings were reduced to 50 principal components, retaining 90.5% of the total variance. These PCA-reduced CLIP features were included as one component of the HighVis feature space. The same CLIP representation was also used to derive the CLIP-predictable component of perceived calorie, as described in the calorie-decomposition section.

*HighVis band construction*

AlexNet, CORnet-S, and CLIP features were grouped into a single HighVis band. This choice was motivated by the goal of constructing a broad visual-semantic baseline rather than adjudicating among individual deep-network architectures. The central scientific question was whether calorie-related predictors explained neural variance beyond general visual and semantic image structure. Therefore, the deep-feature spaces were treated as components of one baseline feature family.

Before concatenation, each HighVis subcomponent was centered and kernel-normalized to unit trace, using the same procedure applied to the LowVis components. This ensured that feature blocks with larger raw variance or dimensionality did not dominate the combined HighVis representation. The kernel-normalized AlexNet, CORnet-S, and CLIP matrices were concatenated and reduced using principal component analysis. The final HighVis band contained 80 principal components and retained 94.9% of the variance of the concatenated high-level feature space. The final HighVis matrix had shape 96 × 80.

*Feature-space overlap diagnostics*

To characterize overlap among feature spaces, representational similarity between feature blocks was quantified using the RV coefficient (Robert & Escoufier, 1976). The RV coefficient is a multivariate measure of similarity between covariance or representational geometries across the same set of observations. Values close to 1 indicate highly similar representational structure across stimuli, whereas values closer to 0 indicate more distinct structure.

Pairwise RV coefficients among high-level feature components ranged from 0.57 to 0.99 (Table S9), indicating substantial representational overlap among the deep-network feature spaces. This supported grouping AlexNet, CORnet-S, and CLIP into a single HighVis band for the primary model family. Importantly, this grouping was not intended to imply that the models are computationally identical, but rather to avoid overinterpreting individual deep-feature spaces when the main hypothesis concerned calorie-related predictors beyond a broad visual-semantic baseline.

*Rationale for dimensionality reduction*

Dimensionality reduction was applied to both LowVis and HighVis to construct compact multivariate feature bands appropriate for the 96-stimulus design. Each outer cross-validation fold contained approximately 77 training images, making extremely high-dimensional feature matrices undesirable for stable model fitting. PCA reduction to 80 dimensions provided a compromise between preserving most of the variance in each feature family and limiting the dimensionality of the feature spaces entering the encoding models.

The dimensionality-reduced feature spaces were not interpreted as isolated neural models of individual feature components. Instead, LowVis and HighVis were used as broad control spaces: LowVis controlled for low-level visual structure, and HighVis controlled for higher-level visual and semantic structure.

**Permutation controls**

Two permutation controls were performed to validate the encoding procedure and to test whether the calorie-related gain was specific to the true ordering of stimuli along the CaloriePredCLIP axis.

The first control was a full condition shuffle. This control tested whether the cross-validation and regularization procedure could produce systematically positive prediction accuracy when the stimulus-response correspondence was destroyed. For each permutation, the 96 condition labels were randomly permuted before constructing the beta matrix, thereby breaking the relationship between image features and neural responses while preserving the remaining structure of the fitting and evaluation procedure. To prevent null models from benefiting from their own optimized hyperparameters, band delta weights were fixed to values estimated in the true model fits, and regularization was set using a scale-relative rule, $\alpha = 100 \times \mathrm{mean}(\mathrm{diag}(K_{\mathrm{train}}))$, without inner cross-validation. This design ensured that positive r_joint values under the permuted null would reflect bias in the cross-validation or fitting structure rather than optimization discovering spurious structure in noise. The control was run for each subject with 100 permutations.

The second control was a calorie-label shuffle. This control tested whether the M2 − M0 gain depended on the specific alignment between food images and CaloriePredCLIP values, rather than on the generic benefit of adding any low-dimensional scalar predictor to the model. Visual features and neural response patterns remained correctly aligned, but the rows of the CaloriePredCLIP feature vector were randomly permuted across images. One hundred shuffled null models were generated per subject. For each subject, a specificity index was computed as the true M2 − M0 gain minus the mean shuffled M2 − M0 gain. Group-level inference on this specificity index was performed using sign-flipping across subjects.

The calorie-label shuffle is conservative because the stimulus set is balanced across high- and low-calorie categories and because CaloriePredCLIP is strongly aligned with broad food-category structure. Some random permutations can therefore partially reconstruct categorical structure by assigning high values to some processed or energy-dense foods and low values to some raw or natural foods. This inflates the shuffled null relative to a fully uninformative baseline. Specificity above this null therefore provides a stringent test that the observed M2 − M0 gain reflects meaningful within-axis alignment rather than merely the addition of a scalar predictor.

**Results**

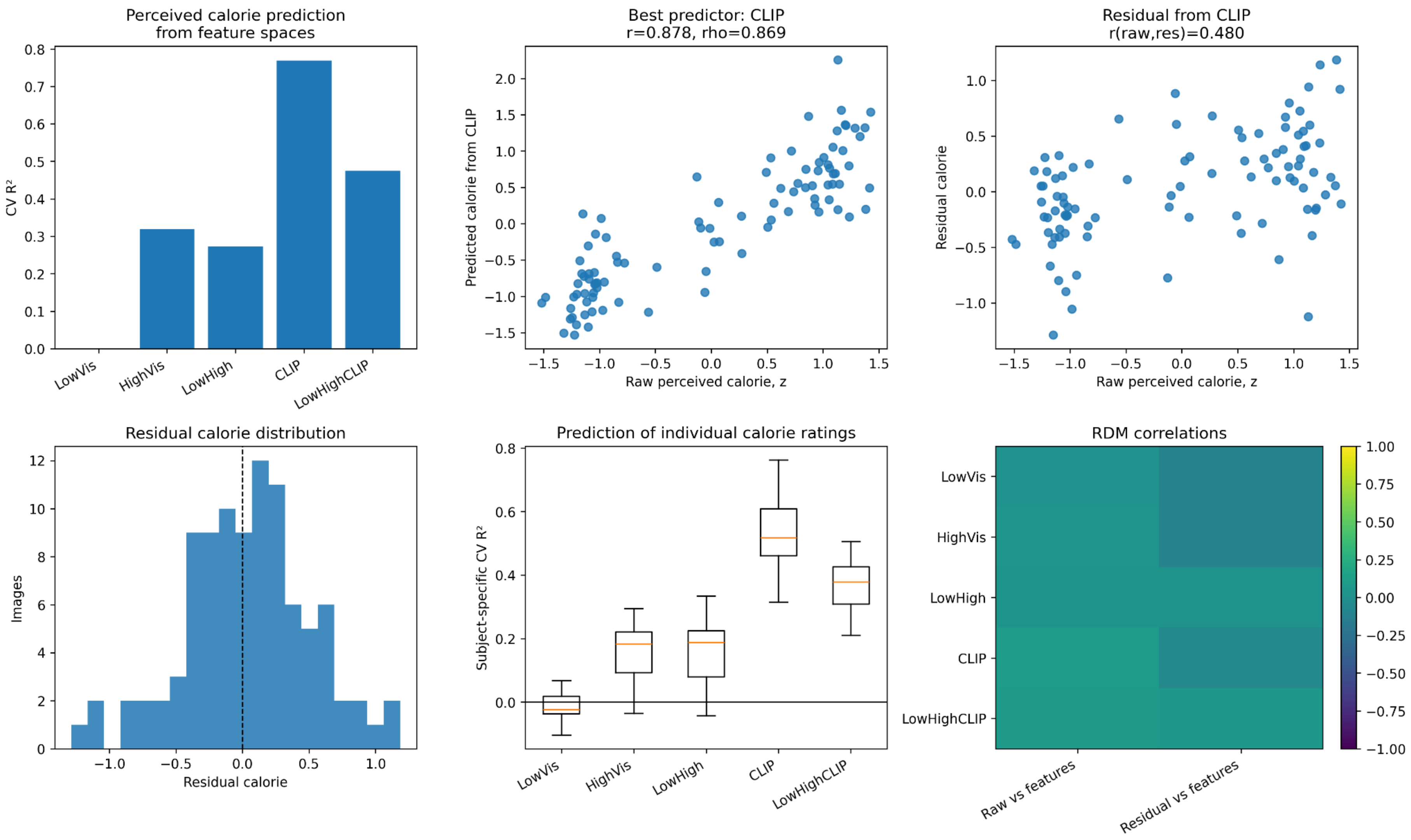


**Supplementary Figure S1. Perceived-calorie prediction diagnostics.**

Cross-validated prediction of perceived calorie ratings from image-feature spaces. The figure summarizes both group-level and subject-specific prediction performance for

LowVis, HighVis, LowHigh, CLIP, and LowHighCLIP feature spaces. Prediction accuracy was quantified using cross-validated explained variance, Pearson correlation, Spearman correlation, and residual variance fraction. CLIP provided the strongest prediction of perceived calorie, indicating that a large proportion of the perceived-calorie axis was recoverable from high-level visual-semantic image *structure. These results motivated the decomposition of perceived calorie into a CLIP-predictable component,*

*CaloriePredCLIP*, and a *CLIP-residual component*, *CalorieResCLIP*.

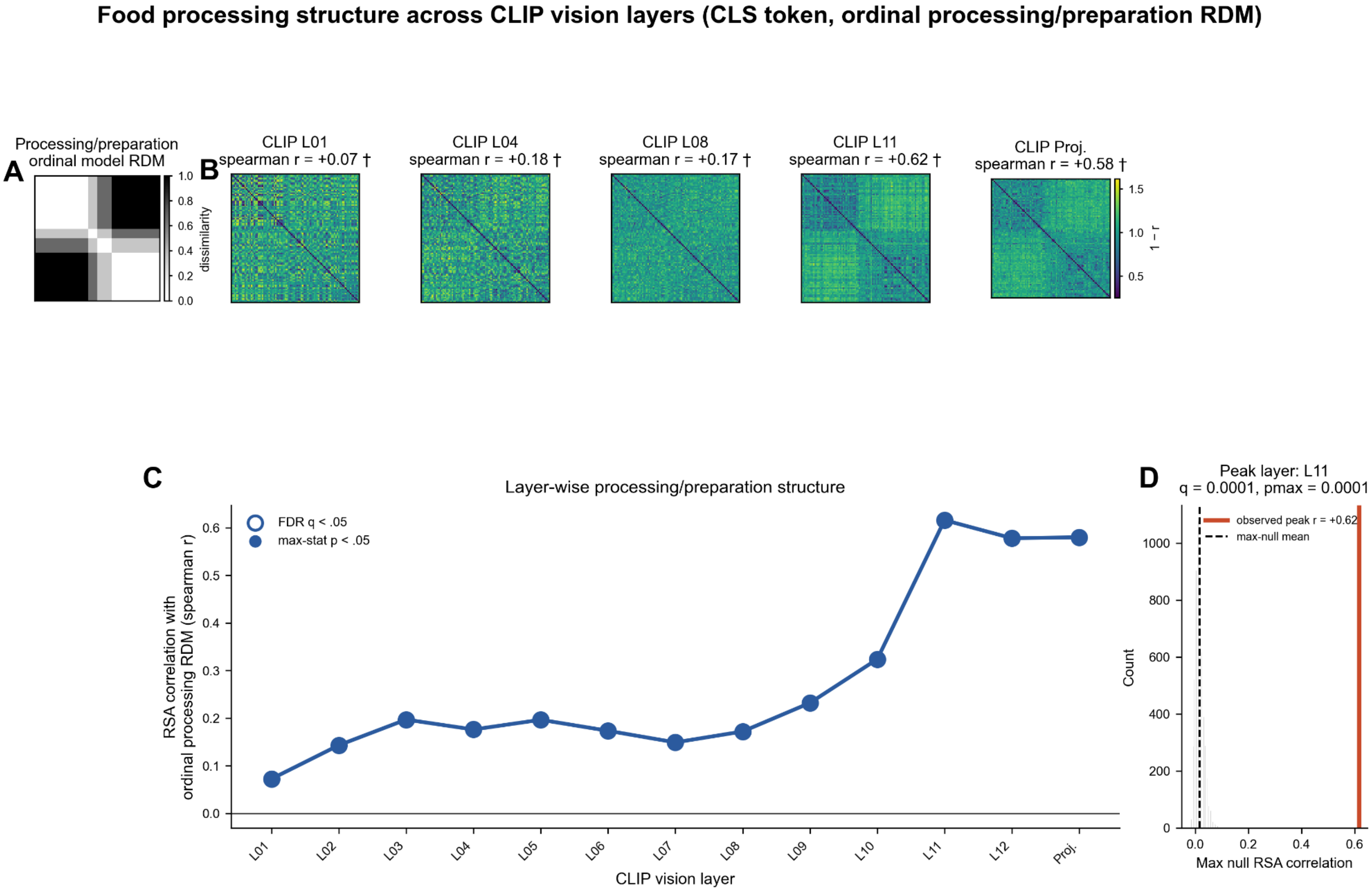


RDMs sorted by processing/preparation level. CLIP RDMs use correlation distance. RSA method: spearman. * = FDR q < .05; † = max-stat familywise p < .05. For CLS-token analyses, the image-independent embedding-stage CLS token is omitted.

**Supplementary Figure S2. Food processing/preparation structure across CLIP vision layers.**

Layerwise characterization of food processing/preparation structure in CLIP visual representations. Processing/preparation-related structure was weak in earlier visual layers and strongest in later CLIP representations and projection space. This pattern indicates that the semantic structure captured by CaloriePredCLIP is not primarily a low-level image property, but instead emerges more strongly in later, higher-level visual-semantic representations.

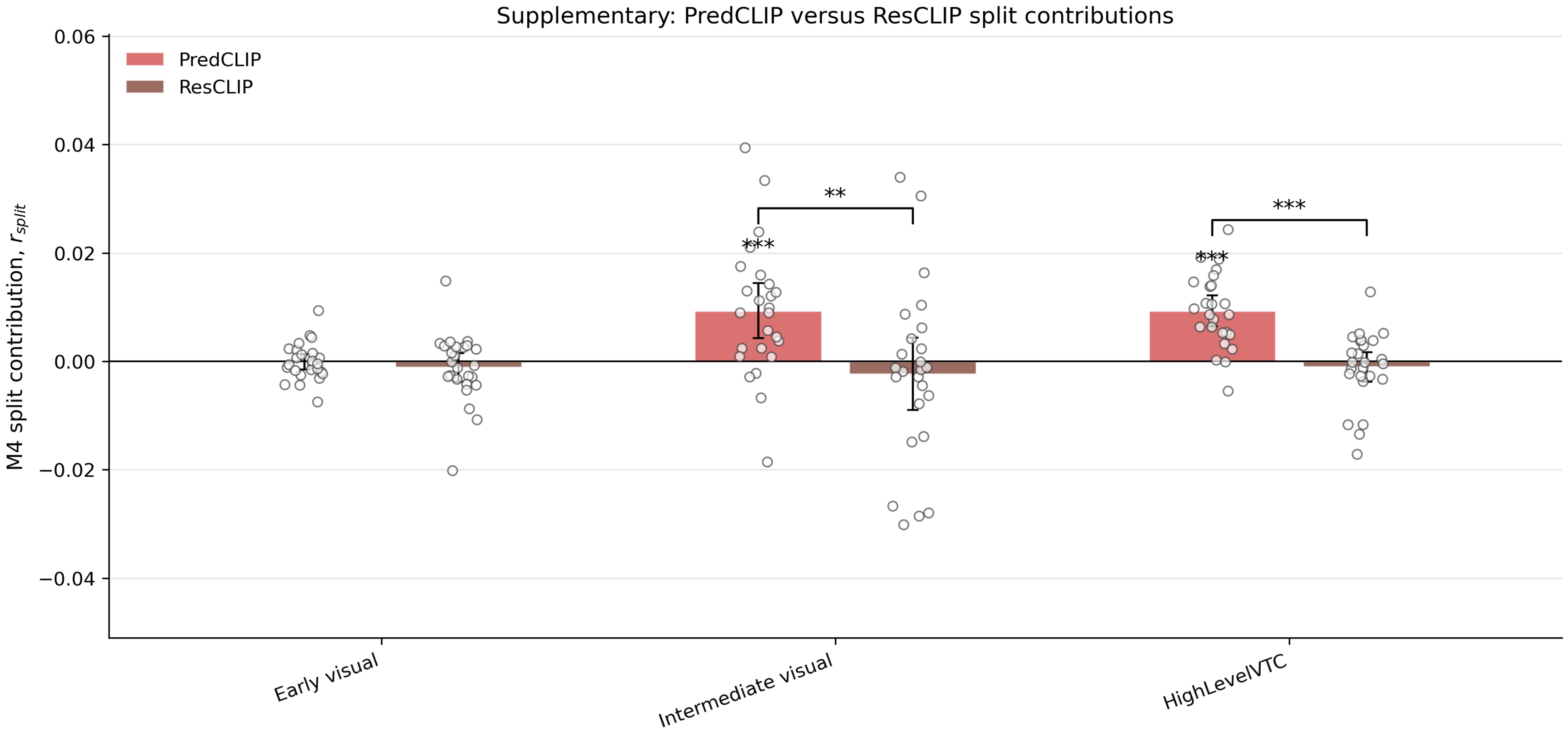


**Supplementary Figure S3. PredCLIP versus ResCLIP split contributions.**

Band-specific split contributions from the full decomposed model, M4, comparing CaloriePredCLIP and CalorieResCLIP across EarlyVisual, IntermediateVisual, and

HighLevelVTC. Split-score analyses showed that CaloriePredCLIP contributed more than CalorieResCLIP in IntermediateVisual cortex and HighLevelVTC, but not in EarlyVisual cortex. These results provide a secondary descriptive decomposition of the joint model and support the main nested-model comparison showing that the CLIP-predictable calorie component was more neurally predictive than the CLIP-residual component in higher-level visual regions.

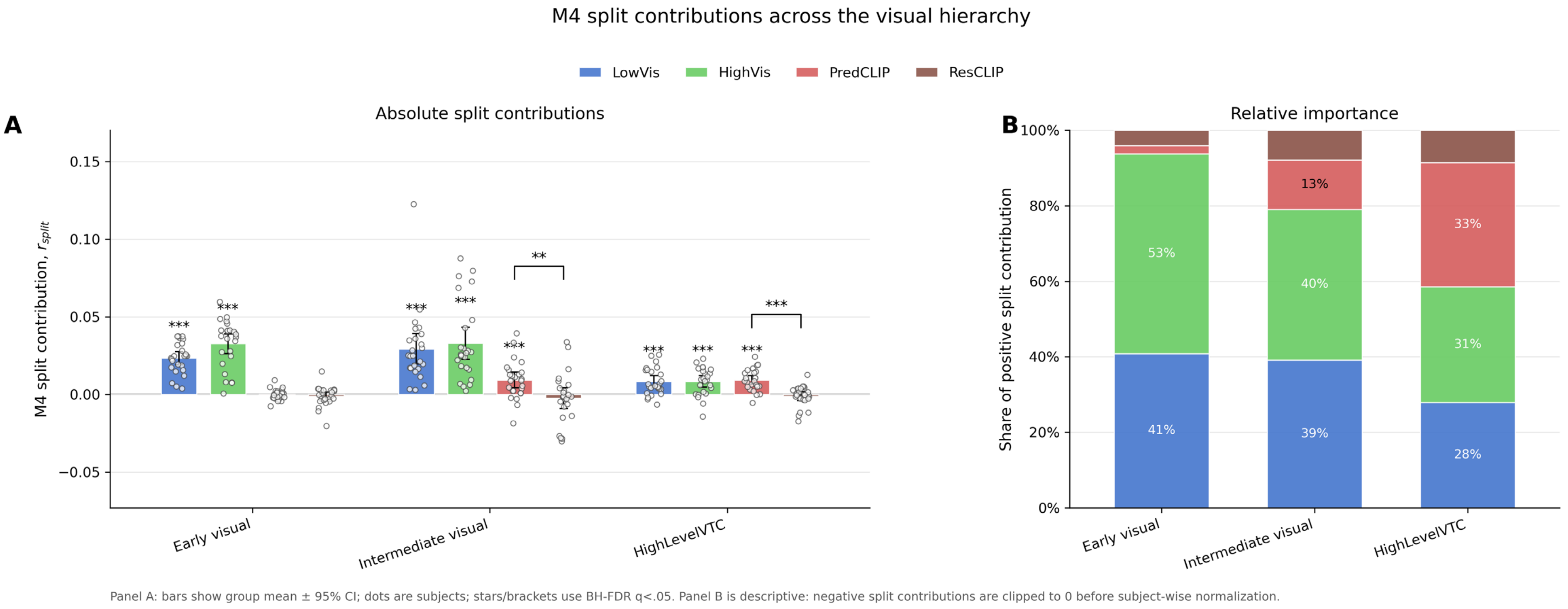


**Supplementary Figure S4. M4 split contributions across the visual hierarchy.**

Absolute and relative split contributions from M4 across the visual hierarchy. The figure shows how LowVis, HighVis, CaloriePredCLIP, and CalorieResCLIP contributed when included in a common model. Across the hierarchy, relative contributions shifted away from generic visual feature spaces and toward CaloriePredCLIP, especially in HighLevelVTC. Split contributions were treated as secondary descriptive measures because they depend on the full model context and on regularization interactions among correlated feature bands.

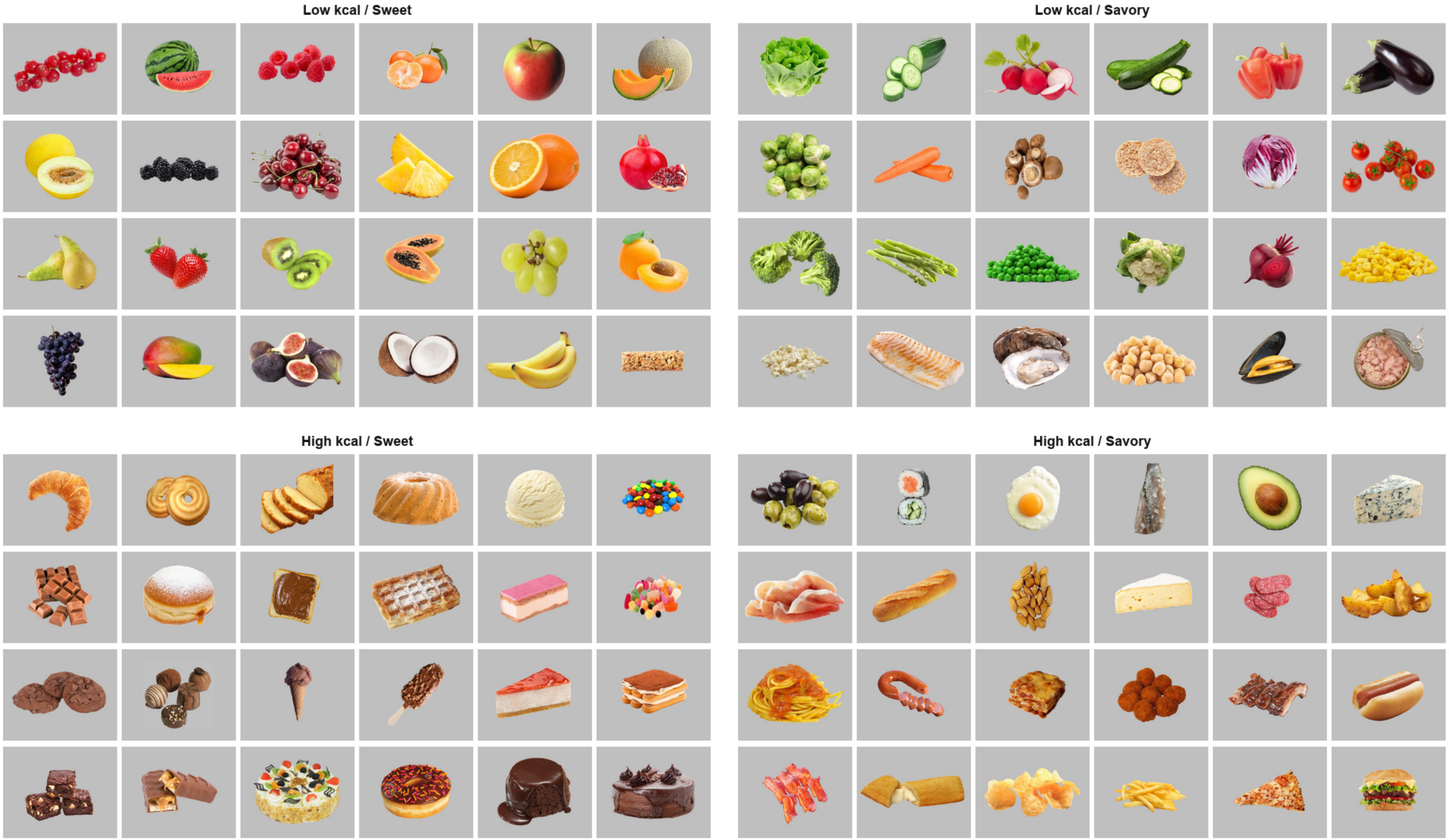
Low kcal / Sweet
Low kcal / Savory
High kcal / Sweet
High kcal / Savory

**Supplementary Figure S5. Full food stimulus set.**

The complete set of 96 food images used in the fMRI experiment, arranged according to the factorial stimulus structure. Images are grouped by perceived calorie category, low versus high calorie, and taste category, sweet versus savory, yielding four balanced stimulus groups: low-calorie sweet, low-calorie savory, high-calorie sweet, and high-calorie savory. All stimuli were presented on a uniform gray background and were used as the image set for the stimulus-level encoding analyses.

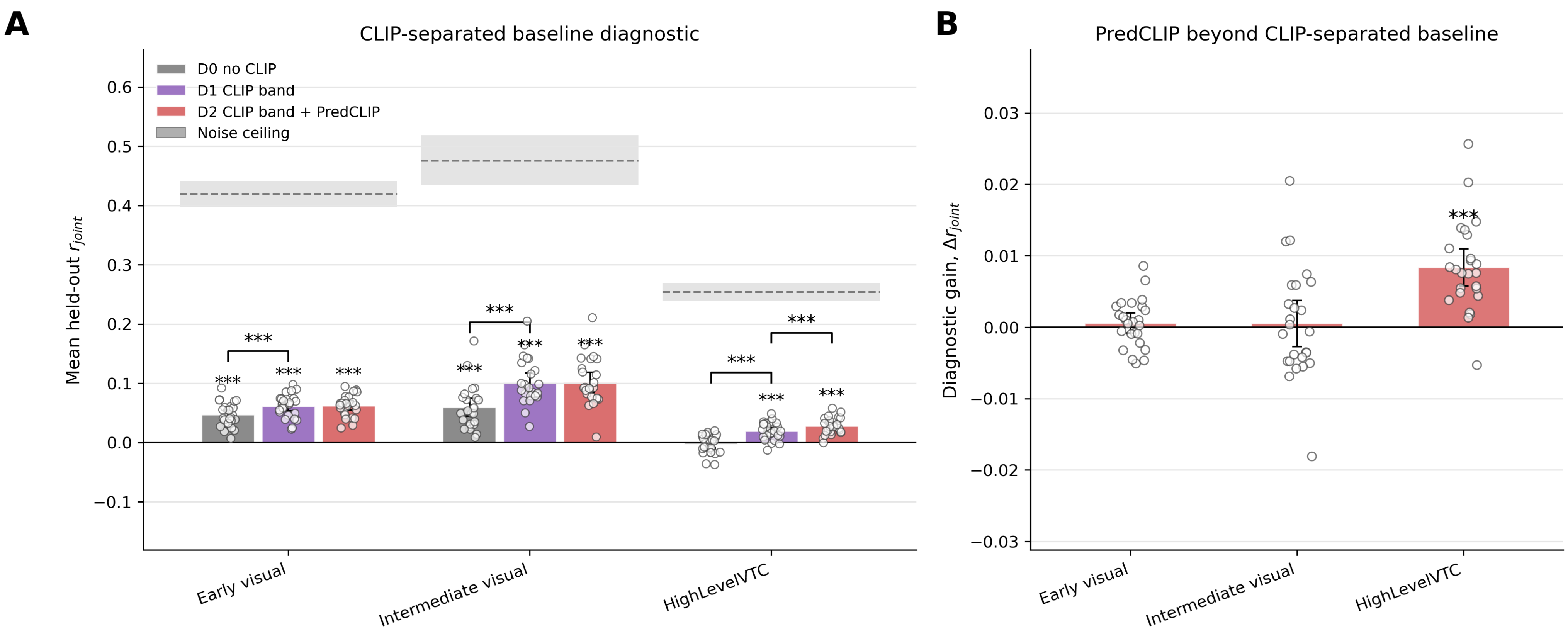


**Supplementary Figure S6. CLIP-separated baseline diagnostic.** (A) Held-out prediction accuracy for the CLIP-separated diagnostic model family across EarlyVisual, IntermediateVisual, and HighLevelVTC. D0 included LowVis and HighVisNoCLIP; D1 additionally included CLIP as a separate feature band; D2 additionally included the CLIP-predicted calorie axis. Bars show group-mean held-out prediction accuracy r_joint, with error bars indicating 95% confidence intervals across participants. Gray horizontal bands indicate split-half noise-ceiling estimates for each ROI, shown as the group mean with 95% confidence interval. Stars above individual bars indicate model performance reliably above zero after BH-FDR correction. Brackets in Panel A are shown deliberately only for the diagnostic steps D1>D0 and D2>D1. The D2>D0 contrasts are omitted from the figure to keep the visualization focused on the effect of interest, namely whether PredCLIP adds beyond a CLIP-separated baseline; the full set of pairwise diagnostic comparisons, including D2>D0, is reported in Supplementary Table S10.

(B) Critical diagnostic gain for PredCLIP beyond the CLIP-separated baseline, computed as D2−D1. Bars show group-mean paired model gain ($\Delta r_{joint}$), with error bars indicating 95% confidence intervals across participants. This comparison tests whether the CLIP-predicted calorie axis explains additional variance beyond a visual baseline in which CLIP already has its own feature band and ridge regularization. PredCLIP produced a reliable additional gain in HighLevelVTC, but not in EarlyVisual or IntermediateVisual cortex.

**Supplementary Table S1. Group-level perceived-calorie prediction diagnostics.**

Note. CV $R^2$ = cross-validated explained variance; residual variance fraction = proportion of perceived-calorie variance not explained by the predictor.

| Predictor | Group CV R2 | Pearson r | Spearman rho | Residual variance fraction |
|---|---|---|---|---|
| LowVis | .000 | .113 | .155 | 1.000 |
| HighVis | .319 | .567 | .585 | .679 |
| LowHigh | .273 | .548 | .543 | .727 |
| CLIP | .769 | .878 | .869 | .229 |
| LowHighCLIP | .475 | .737 | .738 | .525 |

**Supplementary Table S2. Subject-specific perceived-calorie prediction diagnostics.**

Subject-specific perceived calorie ratings were predicted from image-feature spaces using cross-validated ridge regression, and performance metrics were summarized across participants. Values indicate mean ± standard deviation across subjects. CLIP showed the strongest subject-level prediction of perceived calorie, whereas LowVis showed little or negative cross-validated explained variance. These results indicate that the CLIP-predictable component of perceived calorie was not limited to the group-average rating vector but was also present at the level of individual participants' perceived-calorie judgments.

**Note.** CV R² = cross-validated explained variance. Residual variance fraction values greater than 1 can occur when cross-validated prediction performs worse than predicting the training-set mean.

| Predictor | Group CV R2 | Pearson r | Spearman rho | Residual variance fraction |
|---|---|---|---|---|
| LowVis | .000 | .113 | .155 | 1.000 |
| HighVis | .319 | .567 | .585 | .679 |
| LowHigh | .273 | .548 | .543 | .727 |
| CLIP | .769 | .878 | .869 | .229 |
| LowHighCLIP | .475 | .737 | .738 | .525 |

**Supplementary Table S3. Calorie RDM diagnostics.**

Representational diagnostics comparing raw perceived calorie, CLIP-predicted calorie, and CLIP-residual calorie. Correlations are reported between calorie-related representational dissimilarity matrices and feature-space RDMs, as well as among RawCal, PredCal, and ResCal RDMs. RawCal and PredCal showed high representational correspondence, indicating that the CLIP-predicted component preserved most of the representational structure of perceived calorie. In contrast, ResCal showed weak or near-zero correspondence with RawCal and PredCal, confirming that the residual component captured a distinct portion of calorie-rating variance after removing CLIP-predictable structure.

**Note**. RawCal = raw perceived calorie; PredCal = CLIP-predicted calorie; ResCal = CLIP-residual calorie. RDM = representational dissimilarity matrix.

| Predictor | RawCal RDM vs features | PredCal RDM vs features | ResCal RDM vs features | RawCal vs PredCal RDM | RawCal vs ResCal RDM | PredCal vs ResCal RDM |
|---|---|---|---|---|---|---|
| LowVis | .012 | .047 | -.107 | .865 | .035 | -.077 |

| Predictor | RawCal RDM vs features | PredCal RDM vs features | ResCal RDM vs features | RawCal vs PredCal RDM | RawCal vs ResCal RDM | PredCal vs ResCal RDM |
|---|---|---|---|---|---|---|
| HighVis | .043 | .049 | -.109 | .954 | .009 | -.037 |
| LowHigh | .039 | .039 | .027 | 1.000 | .045 | .045 |
| CLIP | .103 | .105 | -.060 | .945 | .015 | -.004 |
| LowHighCLIP | .085 | .085 | .067 | 1.000 | .005 | .005 |

**Supplementary Table S4. ROI-level model performance.**

Held-out prediction accuracy for all nested encoding models across EarlyVisual, IntermediateVisual, and HighLevelVTC. Prediction accuracy is reported as mean r_joint with 95% confidence intervals. Group-level inference was performed across participants using sign-flipping tests on Fisher-z transformed ROI means, with false discovery rate correction applied across the relevant test family. M0 provided reliable prediction in EarlyVisual and IntermediateVisual cortex but not in HighLevelVTC. Calorie-related models improved prediction in HighLevelVTC, supporting the main finding that calorie-aligned predictors contributed beyond the visual-semantic baseline in higher-level visual cortex.

**Note.** r_joint = cross-validated whole-model prediction accuracy, computed as the Pearson correlation between predicted and observed condition-level response patterns for held-out images. p = uncorrected permutation p-value; q = FDR-corrected p-value.

| ROI | Model | N | Mean rjoint [95% CI] | p | q |
|---|---|---|---|---|---|
| EarlyVisual | M0 | 25 | .048 [.037, .058] | < .001 | < .001 |
| EarlyVisual | M1 | 25 | .053 [.044, .062] | < .001 | < .001 |
| EarlyVisual | M2 | 25 | .050 [.041, .060] | < .001 | < .001 |

| ROI | Model | N | Mean rjoint [95% CI] | p | q |
|---|---|---|---|---|---|
| EarlyVisual | M3 | 25 | .050 [.042, .058] | < .001 | < .001 |
| EarlyVisual | M4 | 25 | .049 [.042, .057] | < .001 | < .001 |
| IntermediateVisual | M0 | 25 | .069 [.053, .086] | < .001 | < .001 |
| IntermediateVisual | M1 | 25 | .080 [.064, .097] | < .001 | < .001 |
| IntermediateVisual | M2 | 25 | .076 [.059, .093] | < .001 | < .001 |
| IntermediateVisual | M3 | 25 | .062 [.047, .077] | < .001 | < .001 |
| IntermediateVisual | M4 | 25 | .065 [.049, .082] | < .001 | < .001 |
| HighLevelVTC | M0 | 25 | .002 [-.006, .010] | .283 | .283 |
| HighLevelVTC | M1 | 25 | .021 [.013, .028] | < .001 | < .001 |
| HighLevelVTC | M2 | 25 | .019 [.011, .026] | < .001 | < .001 |
| HighLevelVTC | M3 | 25 | .010 [.004, .016] | < .001 | < .001 |
| HighLevelVTC | M4 | 25 | .018 [.012, .025] | < .001 | < .001 |

**Supplementary Table S4b. ROI-level noise-ceiling estimates.**

Noise-ceiling estimates via repeated split-half for EarlyVisual, IntermediateVisual, and HighLevelVTC. Noise ceilings were available for 21 participants because the implemented split-half procedure required an even number of usable runs; three participants with one excluded run were therefore not included in this descriptive reliability analysis. Noise ceilings were used only as quality-control estimates of reliable stimulus-related signal and were not used for model fitting or primary statistical inference.

**Note.** Values indicate mean split-half reliability estimates with 95% confidence intervals. N = number of participants included in the noise-ceiling analysis.

| ROI | N | Noise ceiling r [95% CI] |
|---|---|---|
| EarlyVisual | 25 | .419 [.398, .441] |
| IntermediateVisual | 25 | .476 [.434, .518] |
| HighLevelVTC | 25 | .254 [.238, .269] |

**Supplementary Table S5. Key encoding model-gain and specificity comparisons.**

Primary model comparisons tested whether calorie-related predictors improved held-out neural prediction beyond the visual-semantic baseline. The main model-gain contrast was M2 > M0, testing whether CaloriePredCLIP improved prediction beyond LowVis + HighVis. The key specificity contrast was M2 > M3, testing whether the CLIP-predictable calorie component explained more neural-predictive information than the CLIP-residual calorie component. M2 improved prediction beyond M0 in all three ROI groups, with the largest gain in HighLevelVTC. M3 did not reliably improve prediction in EarlyVisual or IntermediateVisual cortex, but did improve prediction in HighLevelVTC. Critically, M2 outperformed M3 in IntermediateVisual cortex and HighLevelVTC, indicating that the neural-relevant calorie-related signal was more strongly carried by the CLIP-predictable component than by the CLIP-residual component.

**Note.** Δr_joint values indicate within-subject model differences in held-out prediction accuracy. Confidence intervals are computed across participants. p-values were obtained using sign-flipping tests on Fisher-z transformed within-subject differences; q-values are FDR-corrected.

| ROI | Contrast | Delta rjoint [95% CI] | p | q |
|---|---|---|---|---|
| EarlyVisual | M1 > M0 | .005 [.002, .008] | < .001 | .002 |
| EarlyVisual | M2 > M0 | .003 [.001, .005] | .003 | .008 |
| EarlyVisual | M3 > M0 | .003 [-.002, .007] | .113 | .247 |
| EarlyVisual | M2 > M1 | -.002 [-.004, -.001] | .998 | 1.000 |

| ROI | Contrast | Delta rjoint [95% CI] | p | q |
|---|---|---|---|---|
| EarlyVisual | M2 > M3 | .000 [-.004, .004] | .472 | .871 |
| IntermediateVisual | M1 > M0 | .011 [.006, .017] | < .001 | < .001 |
| IntermediateVisual | M2 > M0 | .007 [.002, .012] | .003 | .008 |
| IntermediateVisual | M3 > M0 | -.008 [-.016, .001] | .969 | 1.000 |
| IntermediateVisual | M2 > M1 | -.004 [-.006, -.002] | 1.000 | 1.000 |
| IntermediateVisual | M2 > M3 | .014 [.005, .023] | .001 | .004 |
| HighLevelVTC | M1 > M0 | .018 [.014, .023] | < .001 | < .001 |
| HighLevelVTC | M2 > M0 | .016 [.013, .020] | < .001 | < .001 |
| HighLevelVTC | M3 > M0 | .008 [.004, .012] | < .001 | .002 |
| HighLevelVTC | M2 > M1 | -.002 [-.003, -.001] | .996 | 1.000 |
| HighLevelVTC | M2 > M3 | .008 [.004, .013] | < .001 | .001 |

**Supplementary Table S5b. Complete model-comparison table.**

Complete set of pre-specified nested model comparisons across EarlyVisual, IntermediateVisual, and HighLevelVTC. This table extends the key comparisons reported in Supplementary Table S5 by including all tested model contrasts, including M4 comparisons. The full comparison table shows that the main conclusions were driven by the improvement of M1 and M2 over M0 and by the advantage of M2 over M3 in higher-level visual ROIs. M4 did not consistently improve over M2, indicating that adding both decomposed calorie components together did not provide substantial additional predictive benefit beyond the CLIP-predictable component alone.

**Note.** M0 = LowVis + HighVis; M1 = M0 + CalorieRaw; M2 = M0 + CaloriePredCLIP; M3 = M0 + CalorieResCLIP; M4 = M0 + CaloriePredCLIP + CalorieResCLIP. $\Delta r\_joint$ values indicate model differences in cross-validated prediction accuracy.

| ROI | Contrast | Delta rjoint [95% CI] | p | q |
|---|---|---|---|---|
| EarlyVisual | M1 > M0 | .005 [.002, .008] | < .001 | .002 |
| EarlyVisual | M2 > M0 | .003 [.001, .005] | .003 | .008 |
| EarlyVisual | M3 > M0 | .003 [-.002, .007] | .113 | .247 |
| EarlyVisual | M4 > M0 | .001 [-.003, .006] | .246 | .493 |
| EarlyVisual | M2 > M1 | -.002 [-.004, -.001] | .998 | 1.000 |
| EarlyVisual | M2 > M3 | .000 [-.004, .004] | .472 | .871 |
| EarlyVisual | M4 > M2 | -.001 [-.005, .002] | .766 | 1.000 |
| EarlyVisual | M4 > M1 | -.003 [-.006, -.000] | .988 | 1.000 |
| IntermediateVisual | M1 > M0 | .011 [.006, .017] | < .001 | < .001 |
| IntermediateVisual | M2 > M0 | .007 [.002, .012] | .003 | .008 |
| IntermediateVisual | M3 > M0 | -.008 [-.016, .001] | .969 | 1.000 |
| IntermediateVisual | M4 > M0 | -.004 [-.012, .004] | .834 | 1.000 |
| IntermediateVisual | M2 > M1 | -.004 [-.006, -.002] | 1.000 | 1.000 |
| IntermediateVisual | M2 > M3 | .014 [.005, .023] | .001 | .004 |
| IntermediateVisual | M4 > M2 | -.011 [-.018, -.004] | .998 | 1.000 |
| IntermediateVisual | M4 > M1 | -.015 [-.021, -.010] | 1.000 | 1.000 |
| HighLevelVTC | M1 > M0 | .018 [.014, .023] | < .001 | < .001 |
| HighLevelVTC | M2 > M0 | .016 [.013, .020] | < .001 | < .001 |
| HighLevelVTC | M3 > M0 | .008 [.004, .012] | < .001 | .002 |

| ROI | Contrast | Delta rjoint [95% CI] | p | q |
|---|---|---|---|---|
| HighLevelVTC | M4 > M0 | .016 [.011, .022] | < .001 | < .001 |
| HighLevelVTC | M2 > M1 | -.002 [-.003, -.001] | .996 | 1.000 |
| HighLevelVTC | M2 > M3 | .008 [.004, .013] | < .001 | .001 |
| HighLevelVTC | M4 > M2 | -.000 [-.004, .003] | .585 | 1.000 |
| HighLevelVTC | M4 > M1 | -.002 [-.005, .001] | .935 | 1.000 |

**Supplementary Table S5c. Hierarchy slope tests.**

Subject-level slope tests assessing whether model gains increased across the visual hierarchy from EarlyVisual to IntermediateVisual to HighLevelVTC. For each participant, model gain was computed within each ROI, and a linear slope was fitted across ordinal ROI levels. Both M2 − M0 and M1 − M0 gains showed significant positive slopes, indicating that calorie-related model gains increased along the posterior-to-anterior visual hierarchy. The primary theoretical test concerned M2 − M0, reflecting the hierarchical increase of the CLIP-predictable calorie component.

**Note.** ROI hierarchy levels were coded as EarlyVisual = 0, IntermediateVisual = 1, and HighLevelVTC = 2. Cohen's dz is the standardized within-subject effect size for the group-level slope test.

| Test | N | Estimate | Cohen dz | p | q |
|---|---|---|---|---|---|
| M2-M0 linear slope | 25 | .007 Delta rjoint per step | 1.29 | < .001 | < .001 |
| M1-M0 linear slope | 25 | .007 Delta rjoint per step | 1.12 | < .001 | < .001 |

**Supplementary Table S5d. M2 − M0 hierarchy pairwise contrasts.**

Pairwise comparisons testing whether the CaloriePredCLIP gain, M2 − M0, differed between adjacent and non-adjacent levels of the visual hierarchy. The M2 − M0 gain was larger in HighLevelVTC than in EarlyVisual and IntermediateVisual cortex, whereas the difference between IntermediateVisual and EarlyVisual did not survive correction. These results indicate that the hierarchical effect was primarily driven by a stronger CaloriePredCLIP gain in HighLevelVTC.

**Note**. Δr_joint values indicate differences in model gain between ROI groups. p-values were obtained using subject-level paired sign-flipping tests; q-values are FDR-corrected.

| Contrast | Delta rjoint | Cohen dz | p | q |
|---|---|---|---|---|
| IntermediateVisual > EarlyVisual | .004 | 0.33 | .057 | .057 |
| HighLevelVTC > EarlyVisual | .014 | 1.29 | < .001 | < .001 |
| HighLevelVTC > IntermediateVisual | .010 | 0.85 | < .001 | < .001 |

**Supplementary Table S6. M4 split contributions.**

Band-specific split contributions from M4, which included LowVis, HighVis, CaloriePredCLIP, and CalorieResCLIP in a common model. Split scores estimate how much each feature band contributed to the jointly fitted prediction and are reported on the Pearson r scale. LowVis and HighVis contributed reliably in EarlyVisual and IntermediateVisual cortex. CaloriePredCLIP contributed reliably in IntermediateVisual cortex and HighLevelVTC, whereas CalorieResCLIP did not show reliable positive split contributions. These secondary analyses support the nested-model results by showing that the CLIP-predictable calorie component made a stable contribution when modeled together with the other feature bands.

**Note.** r_split = band-specific split contribution. Split scores were treated as descriptive because they depend on the full model context and regularization among correlated feature spaces.

| ROI | Band | rsplit [95% CI] | p | q |
|---|---|---|---|---|
| EarlyVisual | LowVis | .023 [.019, .028] | < .001 | < .001 |
| EarlyVisual | HighVis | .033 [.026, .039] | < .001 | < .001 |
| EarlyVisual | PredCLIP | -.000 [-.002, .001] | .549 | .732 |
| EarlyVisual | ResCLIP | -.001 [-.004, .001] | .802 | .802 |
| IntermediateVisual | LowVis | .029 [.020, .039] | < .001 | < .001 |
| IntermediateVisual | HighVis | .033 [.023, .043] | < .001 | < .001 |
| IntermediateVisual | PredCLIP | .009 [.004, .014] | < .001 | < .001 |
| IntermediateVisual | ResCLIP | -.002 [-.009, .004] | .762 | .802 |
| HighLevelVTC | LowVis | .008 [.005, .012] | < .001 | < .001 |
| HighLevelVTC | HighVis | .008 [.005, .012] | < .001 | < .001 |
| HighLevelVTC | PredCLIP | .009 [.006, .012] | < .001 | < .001 |
| HighLevelVTC | ResCLIP | -.001 [-.004, .002] | .776 | .802 |

**Supplementary Table S7. Relative positive split contributions.**

Relative positive split contributions from M4, expressed as percentages within each ROI. Negative split contributions were first clipped to zero and the remaining positive contributions were normalized within participant before averaging across participants. EarlyVisual cortex was dominated by LowVis and HighVis contributions, whereas

HighLevelVTC showed a larger relative contribution from CaloriePredCLIP. These values provide a descriptive summary of how the balance of feature-band contributions shifted across the visual hierarchy.

**Note.** Percentages reflect relative positive contributions and should be interpreted descriptively rather than as inferential model-comparison statistics.

| ROI | HighVis | LowVis | PredCLIP | ResCLIP |
|---|---|---|---|---|
| EarlyVisual | 52.9% | 40.8% | 2.2% | 4.1% |
| IntermediateVisual | 39.9% | 39.1% | 13.1% | 7.9% |
| HighLevelVTC | 30.6% | 27.9% | 32.9% | 8.5% |

**Supplementary Table S7b. PredCLIP > ResCLIP split contrasts.**

Direct comparison between CaloriePredCLIP and CalorieResCLIP split contributions in M4. CaloriePredCLIP exceeded CalorieResCLIP in IntermediateVisual cortex and HighLevelVTC, but not in EarlyVisual cortex. These split-score contrasts provide secondary support for the main specificity result that the CLIP-predictable calorie component carried more neural-predictive information than the residual calorie component in higher-level visual regions.

**Note.** $\Delta r\_split$ = within-subject difference between CaloriePredCLIP and CalorieResCLIP split contributions. p-values were obtained using sign-flipping tests; q-values are FDR-corrected.

| ROI | Contrast | Delta rsplit [95% CI] | p | q |
|---|---|---|---|---|
| EarlyVisual | PredCLIP > ResCLIP | .001 [-.002, .004] | .245 | .245 |
| IntermediateVisual | PredCLIP > ResCLIP | .012 [.004, .020] | .003 | .005 |
| HighLevelVTC | PredCLIP > ResCLIP | .010 [.007, .014] | < .001 | < .001 |

**Supplementary Table S8. Full condition-shuffle control.**

Full condition-shuffle control testing whether the encoding procedure produced positive prediction when the mapping between image features and neural responses was destroyed. For each subject and permutation, condition labels were randomly shuffled before model fitting, thereby breaking the stimulus-response correspondence while preserving the structure of the model-fitting procedure. Null prediction values were close to zero across ROIs and models, indicating that the modeling pipeline did not generate systematically positive prediction under destroyed stimulus-response correspondence. True prediction values exceeded the shuffled null, confirming that the observed model performance depended on the real alignment between image features and neural responses.

**Note.** Null mean ± SD summarizes the distribution of prediction values across condition-shuffle permutations. p(true > null) tests whether the observed model performance exceeded the shuffled null distribution.

| ROI | Quantity | Null mean +/- SD | True mean | p(true > null) |
|---|---|---|---|---|
| EarlyVisual | M0 | -.002 +/- .023 | .074 | < .001 |
| EarlyVisual | M2 | -.002 +/- .022 | .079 | < .001 |
| IntermediateVisual | M0 | -.001 +/- .044 | .093 | .015 |
| IntermediateVisual | M2 | -.001 +/- .043 | .104 | .006 |
| HighLevelVTC | M0 | -.000 +/- .019 | .026 | .083 |
| HighLevelVTC | M2 | -.000 +/- .019 | .030 | .051 |

**Supplementary Table S8b. Calorie-label shuffle specificity.**

Calorie-label shuffle control testing whether the M2 − M0 gain depended on the true alignment between food images and CaloriePredCLIP values. In this control, visual features and neural response patterns remained aligned, but CaloriePredCLIP values were permuted across images. Specificity was defined as the true M2 − M0 gain minus the mean shuffled M2 − M0 gain. Specificity was significantly positive in IntermediateVisual cortex and HighLevelVTC, but not in EarlyVisual cortex, indicating that the CaloriePredCLIP gain in higher-level visual regions depended on the meaningful ordering of images along the calorie-aligned semantic axis rather than merely on adding a scalar predictor.

**Note**. % > 0 indicates the percentage of participants with positive specificity. Cohen's dz is the standardized within-subject effect size for the group-level specificity test. p-values were obtained using sign-flipping tests; q-values are FDR-corrected.

| ROI | N | Mean specificity [95% CI] | Cohen dz | % > 0 | p | q |
|---|---|---|---|---|---|---|
| EarlyVisual | 25 | -.0003 [-.0004, -.0001] | -0.72 | 20.0 | .999 | .999 |
| IntermediateVisual | 25 | .0011 [.0003, .0019] | 0.59 | 72.0 | .001 | .003 |
| HighLevelVTC | 25 | .0010 [.0006, .0014] | 0.99 | 88.0 | .001 | < .001 |

**Supplementary Table S9. RV coefficient matrix for image and behavioral feature spaces.**

RV coefficient matrix quantifying representational overlap among image-feature spaces and behavioral variables across the 96 food images. The RV coefficient ranges from 0 to 1, with higher values indicating more similar representational geometry. Because the RV coefficient is unsigned, values reflect the strength but not the direction of shared structure. Pairwise RV coefficients among the final HighVis components used in the encoding models, AlexNetMid, AlexNetHigh, CORnetIT, and CLIP, ranged from .571 to .721, indicating substantial but incomplete overlap among the deep-network feature spaces. These diagnostics motivated grouping deep-network features into a broad HighVis baseline while preserving CaloriePredCLIP as a targeted, behaviorally interpretable projection of CLIP-readable semantic structure.

**Note.** RV coefficients were used only to characterize overlap among feature spaces and were not used for statistical inference. Because RV is unsigned, high overlap between behavioral variables such as calorie and health indicates shared representational structure but does not specify whether their raw association is positive or negative.

| Feature space | Gabor | Color | AlexNetMid | AlexNetHigh | CORnetIT | CLIP | Palatability | Calorie | Health | Familiarity |
|---|---|---|---|---|---|---|---|---|---|---|
| **Gabor** | 1.000 | 0.149 | 0.395 | 0.333 | 0.404 | 0.329 | 0.035 | 0.070 | 0.084 | 0.040 |
| **Color** | 0.149 | 1.000 | 0.402 | 0.374 | 0.407 | 0.348 | 0.050 | 0.114 | 0.106 | 0.050 |
| **AlexNetMid** | 0.395 | 0.402 | 1.000 | 0.669 | 0.721 | 0.571 | 0.080 | 0.103 | 0.106 | 0.064 |
| **AlexNetHigh** | 0.333 | 0.374 | 0.669 | 1.000 | 0.719 | 0.615 | 0.081 | 0.112 | 0.112 | 0.078 |
| **CORnetIT** | 0.404 | 0.407 | 0.721 | 0.719 | 1.000 | 0.718 | 0.101 | 0.103 | 0.103 | 0.094 |
| **CLIP** | 0.329 | 0.348 | 0.571 | 0.615 | 0.718 | 1.000 | 0.108 | 0.135 | 0.135 | 0.106 |
| **Palatability** | 0.035 | 0.050 | 0.080 | 0.081 | 0.101 | 0.108 | 1.000 | 0.001 | 0.005 | 0.615 |
| **Calorie** | 0.070 | 0.114 | 0.103 | 0.112 | 0.103 | 0.135 | 0.001 | 1.000 | 0.907 | 0.062 |
| **Health** | 0.084 | 0.106 | 0.106 | 0.112 | 0.103 | 0.135 | 0.005 | 0.907 | 1.000 | 0.060 |
| **Familiarity** | 0.040 | 0.050 | 0.064 | 0.078 | 0.094 | 0.106 | 0.615 | 0.062 | 0.060 | 1.000 |

**Supplementary Table S10.** CLIP-separated baseline diagnostic model comparisons. Pairwise comparisons between diagnostic models testing whether the CLIP-predicted calorie axis improves neural prediction beyond a baseline in which CLIP is modeled as a separate feature band. D0 included LowVis and HighVisNoCLIP; D1 additionally included CLIP as an independent feature band; D2 additionally included CaloriePredCLIP. Values are ROI-level held-out prediction accuracies averaged across participants

($r_{joint}$). $\Delta r$ indicates the paired difference between Model A and Model B. Confidence intervals are 95% confidence intervals across participants. Statistical inference was performed using paired sign-flip tests on within-subject Fisher-z-transformed model differences, with BH-FDR correction across the diagnostic comparisons. The D2>D0 contrast is reported for completeness, whereas the critical diagnostic test is D2>D1, which asks whether CaloriePredCLIP explains additional variance beyond the CLIP-separated baseline.

| **ROI** | **Comparison** | **N** | **Model A r** | **Model B r** | **Δr** | **95% CI Δr** | **p** | **q FDR** |
|---|---|---|---|---|---|---|---|---|
| Early visual | +CLIPband > NoCLIP | 25 | 0.0617 | 0.0469 | 0.0149 | [0.0106, 0.0191] | <.001 | <.001 |
| Early visual | +PredCLIP > CLIPband | 25 | 0.0623 | 0.0617 | 0.0006 | [-0.0008, 0.0020] | .195 | .220 |
| Early visual | +CLIPband+PredCLIP > NoCLIP | 25 | 0.0623 | 0.0469 | 0.0155 | [0.0109, 0.0200] | <.001 | <.001 |
| Intermediate visual | +CLIPband > NoCLIP | 25 | 0.1015 | 0.0595 | 0.0420 | [0.0299, 0.0542] | <.001 | <.001 |
| Intermediate visual | +PredCLIP > CLIPband | 25 | 0.1020 | 0.1015 | 0.0005 | [-0.0027, 0.0037] | .365 | .365 |
| Intermediate visual | +CLIPband+PredCLIP > NoCLIP | 25 | 0.1020 | 0.0595 | 0.0426 | [0.0288, 0.0563] | <.001 | <.001 |
| HighLevelVTC | +CLIPband > NoCLIP | 25 | 0.0276 | -0.0004 | 0.0280 | [0.0225, 0.0334] | <.001 | <.001 |
| HighLevelVTC | +PredCLIP > CLIPband | 25 | 0.0355 | 0.0276 | 0.0079 | [0.0053, 0.0105] | <.001 | <.001 |
| HighLevelVTC | +CLIPband+PredCLIP > NoCLIP | 25 | 0.0355 | -0.0004 | 0.0359 | [0.0295, 0.0423] | <.001 | <.001 |

**Supplementary Table S11. Reliability of the CLIP-residual calorie component.**

Split-half reliability of CalorieResCLIP was estimated by dividing rating participants into two independent groups and recomputing the full CLIP-based calorie decomposition separately within each split. Reported values include the split-half Spearman correlation, Spearman–Brown-corrected reliability, the corresponding correlation-scale ceiling, and the $R^2$ ceiling. These diagnostics test whether CalorieResCLIP reflects reliable behavioral structure rather than rating noise.

| Metric | Value |
|---|---|
| Split-half Spearman correlation | .8535 |
| Spearman-Brown reliability | .9206 |
| Correlation-scale ceiling | .9594 |
| R2 ceiling | .9206 |

**Supplementary Table S12. Recovery of CalorieResCLIP from CLIP feature spaces.**

Cross-validated prediction of the final CalorieResCLIP vector from alternative CLIP representations. Models included linear prediction from the original 50-dimensional CLIP representation, nonlinear RBF-kernel prediction from the same space, linear prediction from full 512-dimensional CLIP embeddings, and prediction from discarded higher-order CLIP principal components. Negative or near-zero cross-validated $R^2$ values indicate that CalorieResCLIP was not recoverable from the tested CLIP feature spaces.

| Model | CV R2 | Pearson r | Spearman rho | pperm CV R2 | pperm Spearman | Positive CV R2 fraction of R2 ceiling |
|---|---|---|---|---|---|---|
| CLIP50_linear_ridge | -.0388 | -.2947 | -.3928 | .543 | .995 | .0000 |
| CLIP50_rbf_kernel_ridge | -.0000 | -.1221 | -.2413 | | | .0000 |

| Model | CV R2 | Pearson r | Spearman rho | pperm CV R2 | pperm Spearman | Positive CV R2 fraction of R2 ceiling |
|---|---|---|---|---|---|---|
| CLIP512_linear_ridge | -.0516 | -.1524 | -.0693 | .668 | .358 | .0000 |
| CLIP512_discarded_PC51plus_ridge | -.2000 | -.2096 | -.1863 | .990 | .749 | .0000 |

**Supplementary Table S13. Item-level behavioral and semantic anchors of CalorieResCLIP.**

Item-level correlations between CalorieResCLIP and behavioral, categorical, annotation-based, and CLIP text-axis anchors. Positive correlations indicate stimulus properties associated with higher CalorieResCLIP values, corresponding to foods rated as more caloric than predicted by CLIP. The table reports the strongest anchors by absolute Spearman correlation, with permutation-based p-values and FDR-corrected q-values.

| Anchor | Spearman rho | Pearson r | pperm two-sided | qFDR two-sided |
|---|---|---|---|---|
| predclip_full__PredCLIP_resid_core_annotations_plus_category_z | -.6455 | -.6362 | .0001 | .0003 |
| predclip_full__PredCLIP_resid_core_annotations_z | -.6299 | -.6216 | .0001 | .0003 |
| feature_file__CalorieObjective | .4624 | .4462 | .0001 | .0003 |
| predclip_full__RawCalorie_z | .4609 | .4796 | .0001 | .0003 |
| feature_file__RawCalorie_cvtarget | .4609 | .4796 | .0001 | .0003 |
| feature_file__Calorie_group | .4606 | .4799 | .0001 | .0003 |
| feature_file__Health_group | -.4250 | -.4089 | .0001 | .0003 |
| predclip_full__naturalness_0_3 | -.4249 | -.3625 | .0001 | .0003 |

| Anchor | Spearman rho | Pearson r | pperm two-sided | qFDR two-sided |
|---|---|---|---|---|
| predclip_full__processedness_0_3 | .4246 | .3870 | .0001 | .0003 |
| predclip_full__preparation_0_3 | .4146 | .3815 | .0001 | .0003 |
| predclip_full__raw_produce | -.3927 | -.3384 | .0002 | .0005 |
| predclip_full__fruit_veg | -.3927 | -.3384 | .0002 | .0005 |
| predclip_full__PredCLIP_hat_core_annotations_z | .3866 | .3413 | .0003 | .0007 |
| predclip_full__PredCLIP_hat_core_annotations_plus_category_z | .3733 | .3396 | .0004 | .0009 |
| predclip_full__clip_axis_animal_savory_processed_minus_plant_fresh_z | .3374 | .2796 | .0018 | .0038 |
| predclip_full__clip_axis_energy_dense_treat_minus_light_food_z | .2984 | .2134 | .0041 | .0081 |
| predclip_full__clip_axis_processed_minus_natural_z | .2940 | .2580 | .0050 | .0084 |
| predclip_full__clip_axis_baked_confectionery_minus_fresh_produce_z | .2902 | .2152 | .0043 | .0081 |
| predclip_full__single_ingredient | -.2874 | -.2538 | .0049 | .0084 |
| feature_file__Familiarity_group | -.2568 | -.1652 | .0127 | .0203 |

**Supplementary Table S14. RDM-level anchors of CalorieResCLIP.**

Correlations between a CalorieResCLIP absolute-difference RDM and R1-style behavioral, categorical, and visual model RDMs. These analyses test whether pairwise distances along the CalorieResCLIP axis overlap with pairwise geometry in calorie, health, visual, category, residual-profile, or rating-disagreement model spaces. Permutation inference was performed by shuffling item labels of the CalorieResCLIP vector before RDM construction.

| Anchor RDM | Spearman rho | Pearson r | pperm two-sided | qFDR two-sided | Direct R1 bridge |
|---|---|---|---|---|---|
| Calorie_mean | .1483 | .1694 | .0001 | .0003 | .0000 |
| Color | -.1362 | -.1415 | .0201 | .0536 | .0000 |
| Health_mean | .1209 | .1190 | .0001 | .0003 | .0000 |
| CalorieObjective | .1178 | .1324 | .0001 | .0003 | .0000 |
| Gabor | -.0274 | -.0226 | .5992 | .8055 | .0000 |
| CORnet V4 | -.0208 | -.0257 | .1021 | .2333 | .0000 |
| SavorySweet | -.0106 | -.0094 | .2613 | .4180 | .0000 |
| CORnet IT | -.0065 | -.0219 | .6041 | .8055 | .0000 |
| Palatability_mean | .0024 | -.0158 | .9521 | .9702 | .0000 |
